# A generalized and versatile framework to train and evaluate autoencoders for biological representation learning and beyond: AUTOENCODIX

**DOI:** 10.1101/2024.12.17.628906

**Authors:** Maximilian Joas, Neringa Jurenaite, Dusan Prascevic, Nico Scherf, Jan Ewald

**Affiliations:** Center for Scalable Data Analytics and Artificial Intelligence (ScaDS.AI) Dresden/Leipzig, Leipzig University, Germany; Center for Scalable Data Analytics and Artificial Intelligence (ScaDS.AI) Dresden/Leipzig, Technical University Dresden, Germany; Max Planck Institute for Human Cognitive and Brain Sciences, Leipzig, Germany

## Abstract

Insights and discoveries in complex biological systems, e.g. for personalized medicine, are gained by the combination of large, feature-rich and high-dimensional data with powerful computational methods uncovering patterns and relationships. In recent years, autoencoders, a family of deep learning-based methods for representation learning, are advancing data-driven research due to their variability and non-linear power of multi-modal data integration. Despite their success, current implementations lack standardization, versatility, comparability, and generalizability preventing a broad application. To fill the gap, we present AUTOENCODIX (https://github.com/jan-forest/autoencodix), an open-source framework, designed as a standardized and flexible pipeline for preprocessing, training, and evaluation of autoencoder architectures. These architectures, like ontology-based and cross-modal autoencoders, provide key advantages over traditional methods via explainability of embeddings or the ability to translate across data modalities. We show the value of our framework by its application to data sets from pan-cancer studies (TCGA), single-cell sequencing as well as in combination with imaging. Our studies provide important user-centric insights and recommendations to navigate through architectures, hyperparameters, and important trade-offs in representation learning. Those include reconstruction capability of input data, the quality of embedding for downstream machine learning models, or the reliability of ontology-based embeddings for explainability. In summary, our versatile and generalizable framework allows multi-modal data integration in biomedical research and any other data-driven fields of research. Hence, it can serve as a open-source platform for several major trends and research using autoencoders including architectural improvements, explainability, or training of large-scale pre-trained models.

## BACKGROUND

The key to gaining robust insights into and forming conclusions on complex systems, like cells and organisms, are powerful methods for accurate and informed feature representation of high-dimensional and multi-modal data. This is of particular importance with the onset of increasingly faster and cheaper large-scale data generation studies. However, the availability of multi-modal high-throughput data, such as molecular and genetic (multi-omics) data sets, leads to the curse of dimensionality problem due to a high number of features (e.g. genes, molecules, etc.) in comparison to the sample size (patients, cells, etc.) ^1,2^. Hence, dimensionality reduction methods and unsupervised learning in biology, are increasingly crucial to uncover patterns, discovering biomarkers, enabling efficient supervised learning, or robustly defining similarities of patients which is the key to personalized medicine.

In recent years, deep-learning-based approaches called autoencoders (AE) have become popular for representation learning and (unsupervised) dimension reduction by using a combination of an encoder-decoder structure with a bottleneck layer with low dimension (latent space) ^3^. In comparison to other common approaches like Principle Component Analysis (PCA) or Uniform Manifold Approximation and Projection (UMAP), AEs and their variants potentially offer advantages, such as non-linear and deterministic embeddings, flexible multi-modal data integration, or even generation of synthetic data ^3,4^. The latest variants of proposed AE architectures are even capable of data modality translation (cross-modal AEs) ^5^ or incorporating domain knowledge to gain explainability of the latent space and embedding ^6–8^.

These strong points of AE are particularly helpful and have become popular for multi-modal data integration in biology due to the complex interactions of genetic and molecular factors in cells, tissues, and whole organisms ^9–15^. Further, single omics measurements only shed light on one layer of these interactions, such as gene expression (transcriptomic), and cannot fully capture the origins of a disease (e.g. cancer), which are multi-factorial and perturb cell processes across all layers including epigenetic, genomic, metabolomic or proteomic cellular states ^16^. To this end, a data set like the "The Cancer Genome Atlas (TCGA)" is immensely valuable as it contains multi-omic measurements of thousands of patients across all major cancer types ^17^. However, developing methods to integrate all data modalities and shed light on the complex interplay via low dimensional representations is still a major challenge ^18^.

Despite the advantages and active research on AE architectures and variants, their widespread application in biological representation learning and other domains is hampered by a lack of standardized and generalized frameworks for user-friendly and efficient training of various AEs. Currently, there is no single framework that combines several key aspects such as state-of-the-art AE variants, simultaneous integration of multiple data modalities, and flexible configuration as well as hyperparameter optimization. A recently published framework, Rapidae ^19^, offers some aspects, but has a special focus on time series data and does not offer a full pipeline for (biological) representation learning including data preprocessing, visualizations, and evaluation of embeddings.

To this end, we developed the PyTorch-based ^20^ open-source framework, AUTOENCODIX, which combines these aspects for widespread application for biological representation learning with AEs. While primarily focused on large-scale molecular genetic data, AUTOENCODIX’s flexible and generalized design enables its use in various other fields. Having a unified implementation of AEs and easy parameterization at hand, we provide in this study a first wider benchmark of AE variants for common tasks in biological representation learning, including embedding performance for prognostic clinical models, data modality translation, and explainability by design with ontology-based AEs.

## DESIGN AND FEATURES

Our Python and PyTorch-based framework was developed to enable the applicability of AE architectures for a broad range of multiomics data integration tasks and beyond (*cf.* Fig. 1). Hence, AUTOENCODIX resolves the limitations of current frameworks and AE implementations by: (1) introducing multiple AE architectures in (2) one end-to-end pipeline, from preprocessing to downstream analyses, such as, visualizations and evaluation of embeddings; supporting (3) multi-modal data integration; (4) hyperparameter tuning; (5) flexible and explainable usability for data-driven biomedical research and, lastly, (6) easy extensibility to new AE architectures and methods.

**Figure 1:**
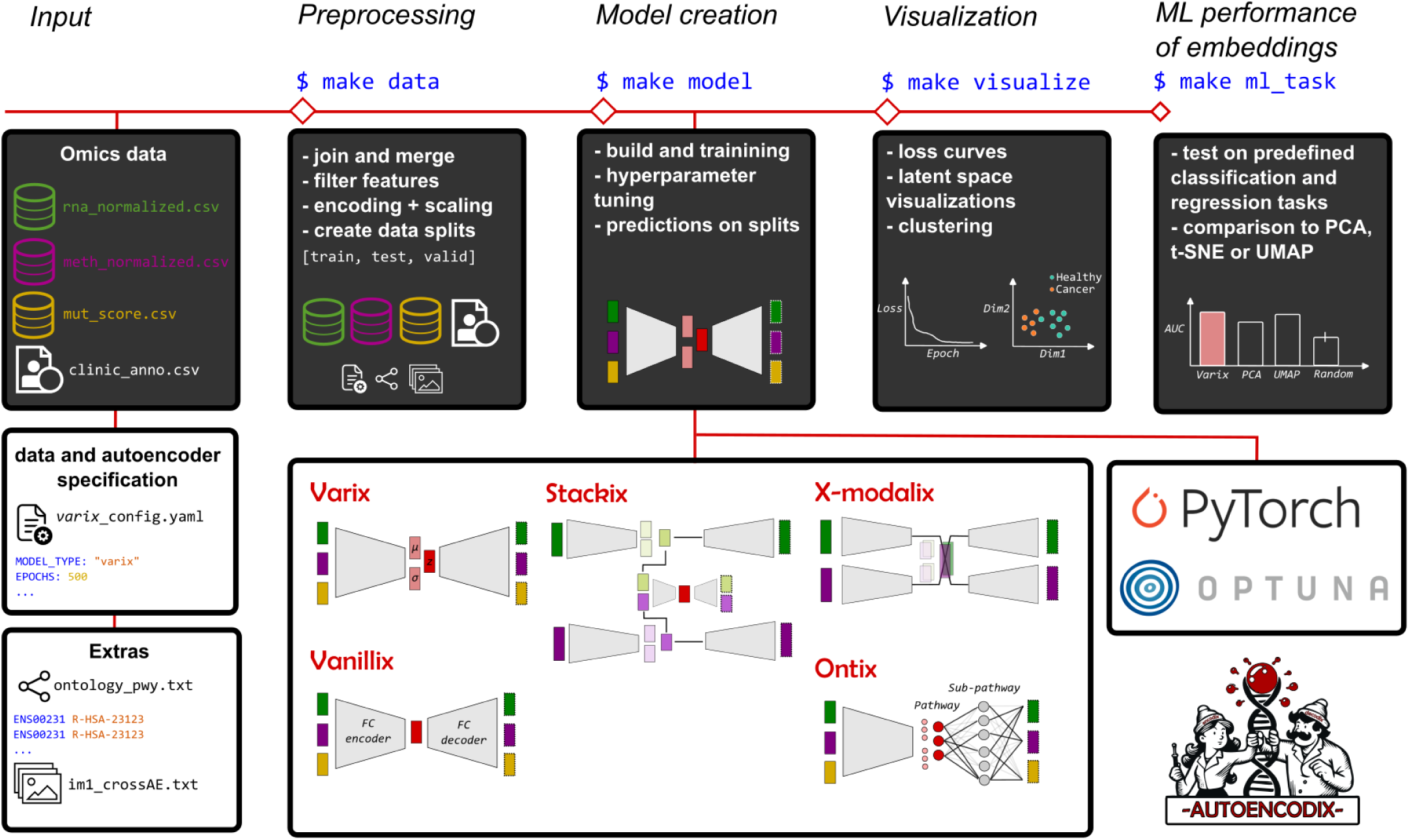
Schematic illustration of open-source framework, AUTOENCODIX, as a full pipeline for training and evaluation of various AE architectures. Upon release, five common or emerging AE architectures are available and shown as sketches. See sections Methods: Framework and autoencoder implementation details, and Results: Design and features, for a detailed description.

As input, AUTOENCODIX can handle any number of data modalities where each data modality, including biological annotations, is entered as a data frame. While we offer scaling of input data modalities with several methods, we assume that each data modality is normalized beforehand for any technical bias (e.g. count normalization by library size etc.). For single-cell data integration, we include preprocessing steps for the AnnData ^21^ h5ad-files, allowing for the extraction of relevant data modalities and annotations for AE training. All options and data input are defined via a YAML configuration file for easy parameter benchmarking and user-friendly access to a modular framework while maintaining full control over the pipeline. In addition, configuration files ensure transparent and reproducible data analyses, enabling openness by design.

Each of the following steps in our pipeline can be either conducted individually or in a single command: data preprocessing, model creation and training, visualization, and evaluation of embeddings on user-defined machine learning tasks (e.g. cancer subtype classification or survival prediction) based on annotation data of samples (labels). A full list and description of options and features for the preprocessing step, such as feature filtering or model hyperparameters, can be found in the AUTOENCODIX documentation).

At the core of AUTOENCODIX are five different AE architectures for a wide range of applications, which capture many of the recent developments in model and method research of the field. Firstly, we present two baseline models, a vanilla autoencoder (Vanillix) and a variational autoencoder (Varix), with fully connected encoder and decoder layers and a bottle-neck layer to produce the latent space. All fully-connected layers are composed of a series of linear, batch normalization, dropout, and ReLU activation layers. Secondly, we implemented a stacked (Stackix) variational autoencoder (also called hierarchical autoencoder), which shows good performance for multi-omics data integration ^22^. It can handle multiple data modalities by learning individual VAEs first, and each latent space is then combined into a concatenated input for a stacked VAE to integrate the data modalities. Explainability in VAEs is still an active field of research, in particular for biomedical applications, however here we implemented a robust approach to incorporate biological knowledge, such as molecular pathways, into the decoder connectivity of VAE, as shown by ^6–8^. Next, our implementation of the ontology-based VAE, Ontix, can handle either a single/linear or two-layer hierarchy of features and provides explainability to latent dimensions because they represent e.g. biological pathways or chromosomes. Lastly, recent and exciting progress has been made in using VAEs to translate between different data modalities of multi-omics and imaging data by Yang et al. ^23^, which is referred to as the cross-modal VAE, X-modalix here. We also support the translation between two data modalities, including image data, while other architectures are limited to numerical and categorical features.

Apart from its features, AUTOENCODIX is unique to other frameworks and packages in our commitment to long-term maintenance of the codebase, the future development to incorporate new trends, as well as providing a contribution guide for the community. This is often lacking in other implementations hindering adoption by the community. Taken together, these points ensure the sustainability of our open-source framework, which can serve as a platform for the community to facilitate research on autoencoders, not only in biological representation learning.

Another key shortcoming of other implementations and frameworks is that hyperparameter tuning, visualizations and embedding evaluation are not supported. In AUTOENCODIX, tuning is in-built because it is critical for real-world applications in biological representation learning and beyond. In the following, we will demonstrate most of the described features using multiple experiments, benchmarks, and showcases on publicly available data sets providing both a user-centric illustration of the capabilities of our framework as well as important new insights into aspects of AEs for biological representation learning.

## RESULTS

### Navigating the zoo of autoencoders for representation learning

In our first set of experiments, using two multi-omics data sets, the pan-Cancer Genome Atlas (TCGA) and a single-cell data set of cortical development (see Data sets), we demonstrate the model performance of the AUTOENCODIX architectures; Vanillix, Varix, Ontix and Stackix (see Fig. 2A). By performing tests on architectural choices, data modalities, default hyperparameters and their tuning, we aim to provide user recommendations as well as practical insights into the characteristics of AEs. In the next sections, and via a second set of experiments, to demonstrate wide-range medical applicability, we explore the explainability and cross-modal data translation directions of our models.

**Figure 2:**
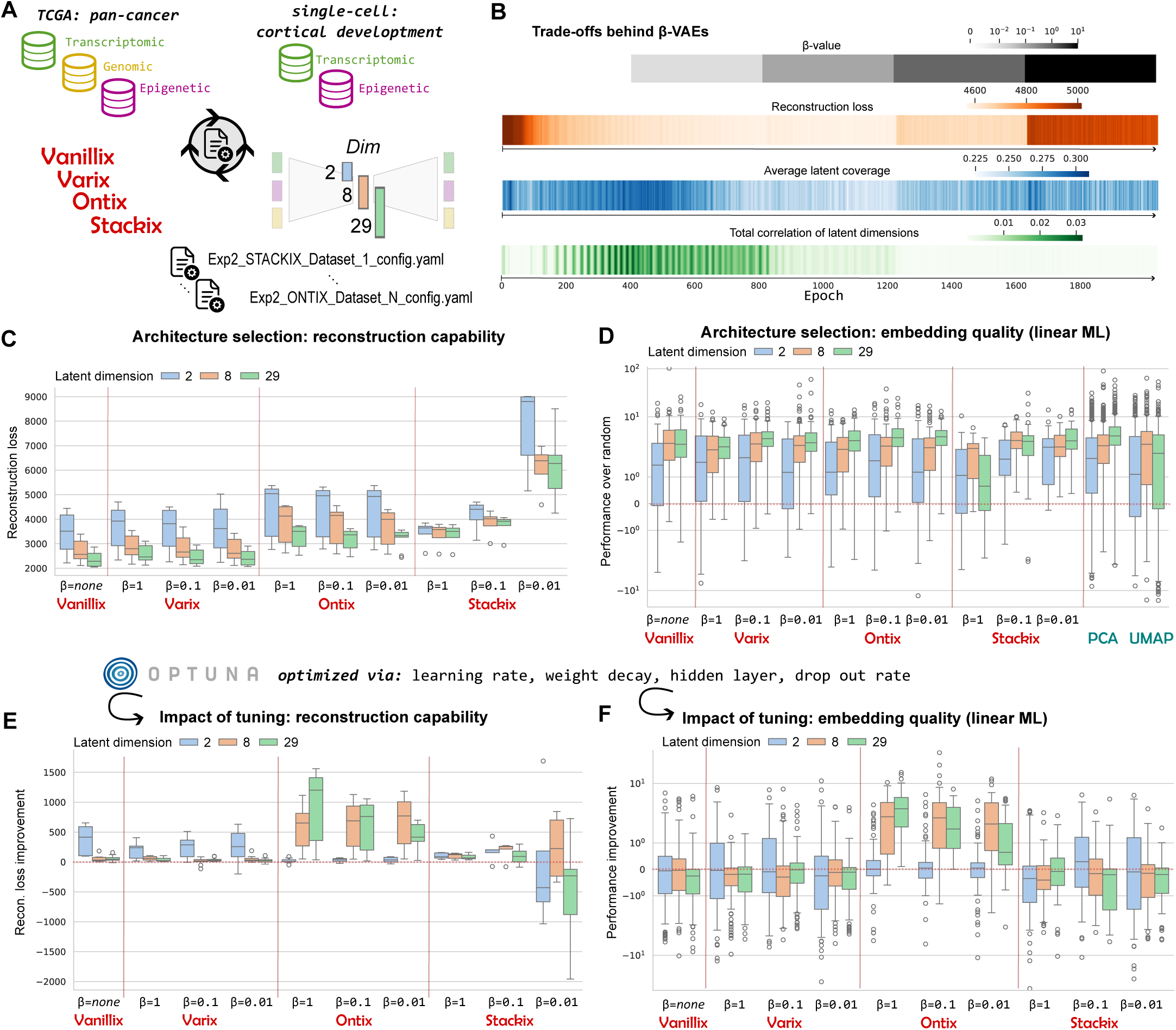
AUTONENCODIX and its architectures for representation learning. **(A)** Overview of the data sets used, their Omics modalities, and AE architectures for comparison of performance. **(B)** Illustration of trade-offs behind *β* configuration in VAEs via training of two-dimensional Varix on TCGA data set and all data modalities with altered *β* every 400 epochs. **(C)** Aggregated reconstruction loss of all combinations as shown in (A) of fully tuned AEs and in comparison to non-tuned AEs in **(E)**. **(D)** Aggregated results of embedding performance of each approach on multiple downstream machine learning tasks (via linear regression or logistic regression), shown as normalized performance over randomly selected features with a similar dimension to the AE. **(F)** shows embedding performance in comparison to non-tuned AEs. Red dotted lines in (D), (E) and (F) indicate the boundary (zero) between improvement (positive-values) and worsening of the performance (negative-values).

#### Reconstruction is not all you need: trade-offs behind *β*-VAE

Variational autoencoders (VAE) are probabilistic generative latent models, where the latent space is modelled as a distribution (typically a multivariate normal). To train VAEs, the encoder approximates this latent (posterior) distribution and the loss function is extended by a measure of similarity of the estimated latent distribution with the prior using, e.g. Kullback–Leibler divergence (KL loss). Hence, VAEs are trained for a trade-off between minimizing the reconstruction loss while encoding and decoding samples, and the compactness of the latent space. The trade-off is captured by the weighting factor, *β*, between the loss terms (see Autoencoder architectures and implementation - general) and we analyzed the impact of *β* on the latent space by using an annealing strategy, where *β* is increased every 400 epochs in five steps from zero to ten. Since all architectures, besides Vanillix, are VAEs and *β* is a critical hyperparameter and not directly tunable via loss function optimization, its configuration is a necessary preceding step for most use cases.

By step-wise increasing *β*, we can analyze the impact of KL-loss on the reconstruction capability, the latent space density (average coverage per latent dimension), and the total correlation of latent dimension, a measure of independence of latent dimensions (disentanglement, cf. PCA) ^24^. Reconstruction loss is as expected the lowest for very small *β* = 0.01 (see Fig. 2B). This behaviour is desirable when the low-dimensional representation must contain the maximum amount of information on high-dimensional Omics data. However, larger weights of KL-loss result in latent dimensions with higher density, and in addition, higher disentanglement, which is advantageous for representation learning e.g. to get disentangled (independent) and more interpretable dimensions. Additionally, when using VAEs to generate realistic synthetic data, a connected and dense latent space is preferable in order to generate meaningful synthetic high-dimensional data from any point in the latent space.

Based on these results, we will use the three *β*-values (0.01, 0.1 and 1) with reasonable trade-offs for further testing of VAE architectures in comparison to the vanilla AE.

#### AUTOENCODIX enables architecture comparison and hyperparameter analysis

One of the main motivations for the AUTOENCODIX implementation is the ability to easily compare different types of AEs and associated hyperparameter configurations to find the best combination for a given data set or task. Although there have been some benchmarks ^11,22^, there are little to no recommendations regarding the strengths and weaknesses of different architectures, the impact of tuning, or the performance of embeddings in downstream machine learning tasks.

Here, as a first step, we analyzed all four types of AE, on two data sets (TCGA and single-cell) with up to three different data modalities and three latent dimensions (see Fig. 2A and Experiments). We chose the three levels of latent dimensionality to represent typical use cases; (*L_dim_* = 2), a two-dimensional visualization, (*L_dim_* = 8), an intermediate dimensionality, and (*L_dim_* = 29), motivated by prior knowledge on biological ontology (number of Reactome top-level pathways). Further, using in-built Optuna support ^25^, we tuned hyperparameters for learning rate, weight decay, hidden layer structure, and dropout rate. To show the effect of tuning, we also determined the performance of the respective standard AE configuration.

AEs are often used to determine low-dimensional representations (embeddings) to enable better training of supervised learning models, which is not directly captured by the loss function optimization. Hence, we consider two performance criteria; reconstruction capability, i.e. reconstruction loss, (see Fig. 2C and cf. Supplementary Figure S1 for non-aggregated results) and evaluated the embedding performance on downstream regression and classification tasks using three common ML algorithms (see Fig. 2D for performance on linear ML and cf. Supplementary Figure S2 for all three). Tasks are based on available sample annotations (a full list is given in the Supplementary Table S3) and include e.g. cell type and cancer subtype classification or prediction of survival duration (regression). As a baseline, we determined embedding performance as the normalized improvement over training on randomly selected features (matching the number of latent dimensions). We also compared the embeddings with PCA and UMAP for cross-method validation.

As expected, we observe that the reconstruction capability of AEs with no KL-loss (Vanillix) or low *β* are more precise in recovering input from the latent space (see Fig. 2C). Similarly, the smaller the latent space, the lower the reconstruction capability, where, in particular, two-dimensional VAEs struggle to precisely recover high-dimensional input data. Both results and general trade-offs in reconstruction capability are largely independent concerning data modality, data set, and VAE type.

The stacked VAE (Stackix, see also Methods - Stacked variational autoencoder) demonstrates similar performance, but is not directly comparable to other architectures due to its uncoupled two-phase training process of each VAE and final reconstruction loss is calculated with embeddings as input from each data modality, rather than on features as in other AE types. Fig. 2C shows the sum of reconstruction loss of individual VAEs per data modality and the final stacked VAE. To this end, the reconstruction loss of Stackix is generally higher compared to other architectures. A lower *β*-configuration has a higher impact in Stackix, as the embeddings of individual VAEs will have a larger data range leading to a higher MSE-loss. Notably, the ontology-based VAE (Ontix, see also Methods - Ontology-based variational autoencoder) demonstrates diminished reconstruction ability, which is justified by the constraints of its sparse decoder reflecting the gene involvements in Reactome pathways.

Compared to non-tuned AEs, reconstruction performance improves primarily for two-dimensional vanilla AE and VAE architectures, while tuning hyperparameters leads to an increase in reconstruction loss for the stacked VAE and *β* = 0.01 (see Fig. 2E). This occurs because only the sub-VAEs of the stacked VAE can be tuned individually, not in their final combined form. Hence, tuned sub-VAE’s improve individually with a more detailed embedding, but larger range of latent dimensions, which leads again to higher reconstruction loss. On the other end, ontology-based VAEs with latent dimension 8 and 29 benefit the most from tuning, while other architectures are mostly comparable in performance when using standard configurations.

#### Reconstruction capability and embedding quality are not directly correlated

Besides the reconstruction capability of AEs, the performance of embeddings for downstream learning of prognostic machine learning models is crucial for application in biological representation learning. In our experiments across two data sets, all data modality combinations, and three machine learning algorithms, no single AE architecture consistently outperforms the others; results vary based on the combination of architecture and algorithm used (see Fig. 2D, using linear ML algorithms, cf. Supplementary Figure S2). Including other methods (PCA and UMAP) and using random features as a reference, we can show that all dimension reduction methods outperform random feature selection not only on average, but on nearly all tasks. Further, we obtained insight that linear ML algorithms benefit in their performance over randomly selected features, but, interestingly, RandomForest and SVL (radial kernel) show inverse behavior (see Supplementary Figure S2). From the user-perspective, we conclude that the evaluation of the embedding performance for downstream tasks is crucial and all architectures should compared in practice, since there is no general superiority of a single architecture or an established method like PCA or UMAP. This is in particular true for VAE with higher *β*-values or ontology-based VAEs where we can see that worse reconstruction capability does not correlate with embedding quality for prognostic models. The same applies when analyzing the impact of tuning on embedding quality. Except for ontology-based VAEs with latent dimensions of 8 and 29, the embeddings show no consistent improvement (see Fig. 2F). This underscores that improvements in reconstruction loss do not necessarily result in better embeddings. However, large improvements in reconstruction as for some ontology-based VAEs and others, will likely significantly improve the performance of embeddings for prognostic model training.

On the other hand, it seems that our standard configuration of AEs is fairly robust for a broad range of applications without any hyperparameter tuning. An exception is the ontology-based VAE which has, in general, different hyperparameter requirements due to their sparse decoder structure. For example, they show a necessity for lower dropout rates and faster learning rates compared to other models (see Supplementary Figure S3), which suggests alterations to our standard configuration for this type. We will analyze in the next section that these parameters might heavily influence the reliability of latent space explainability.

### Ontix provides explainability of latent space by-design

#### Ontology is the cornerstone to gaining biological insights

A key advantage of AEs over methods like UMAP is their deterministic embedding generated by the encoder structure. Secondly and importantly, they allow for greater explainability of the latent space dimensions by incorporating (biological) knowledge. The latter is achieved by restricting the decoder to feature connectivity according to an ontology (see Fig. 3A) which creates explainability ‘by-design’, rather than the ‘post-hoc’-like feature contribution or importance scores.

**Figure 3:**
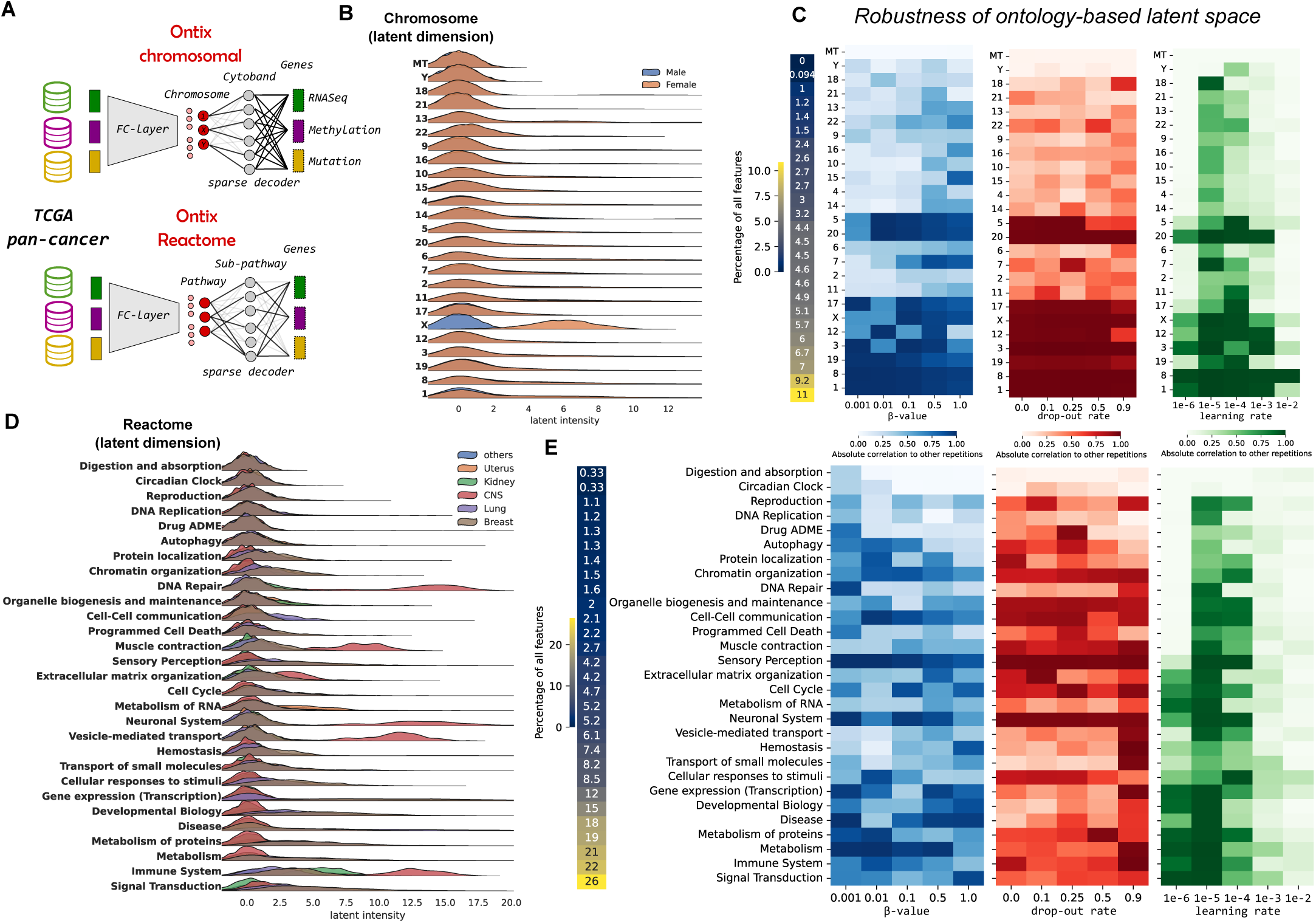
Ontix: biologically-informed decoder enables explainability of the latent space, but its robustness depends on the hyperparameters. **(A)** Two types of ontologies are tested on the TCGA data set: chromosomal-based and Reactome-pathway-based latent dimensions. **(B)** and **(D)** show ridge-line plots of latent intensity distributions, based on selected examples of sample classes (sex and cancer tissue of origin) to show applicability for explainability. **(C)** and **(E)** show the robustness of these embeddings depending on the hyperparameters. Robustness is defined as the mean absolute Pearson correlation between five independent training runs with a randomized data split and weight initialization.

We tested our implementation again on the TCGA (see Fig. 3) and the single-cell cortex data sets (see Supplementary Figure S4) with two types of ontologies: chromosomal location of genes (features) and biological pathways (Reactome database ^26^). Chromosomal location as ontology is ideal for testing and benchmarking since there is no overlap between the features and ontologies, as in biological pathways, and we have a testable expectation for a signal on sex-related chromosomes, separating male and female samples. Indeed, we can observe a strong signal in the latent dimension, representing the X chromosome for TCGA cancer samples (see Fig. 3B). Interestingly, there is no signal on the Y chromosome for TCGA which is related to the small size of the chromosome and very few genes (features) supporting this latent space dimension in this data set. Without the incorporation of an additional loss function weighting to balance connectivity, some ontology-dimensions will be disregarded in favor of capturing the majority of the data variance.

A realistic application of an explainable latent dimension is given in Fig. 3D, where we show latent intensities associated with the biological pathways depending on the tissue of cancer origin. Based on this, one could conclude that cancers of the central nervous system are very distinct from other cancer types, where its characteristic is related to biological processes such as the immune system, vesicle-mediated transport or DNA repair.

#### Getting the hyperparameterization right for robustness

A previously raised issue with biologically-informed neural networks is that they might not be robust to randomness in training ^27,28^, which likely affects ontology-based VAEs. To study robustness, we calculated the correlation of latent spaces across five repetitions, including randomized train-test-valid splits, weight initialization, etc., and varied three key hyperparameters for ontology-based VAEs: weighting factor *β*, dropout rate (encoder layers), and learning rate during training (see Fig. 3C and 3E).

We selected hyperparameters based on the following expectations: (a) *β* limits the flexibility of the latent space, (b) the dropout rate mitigates instability caused by gene-to-pathway overlaps, and (c) the learning rate impacts training stability. Based on our results we observe generally that robustness across ontology-terms is highly non-uniform and dependent on the combination of signal and the number of genes they are connected to. In particular, our chromosome-based Ontix shows that larger chromosomes are more robust (see Fig. 3C using TCGA data and cf. Supplementary Figure S4 using single-cell data). Further, we see that the learning rate has the overall biggest influence on robustness and should be set slower than previous hyperparameter tuning results suggest. This is critical as it demonstrates a trade-off between loss function optimization and the reliability of latent space-derived biological insights. On the other hand, *β*-value and dropout rate do not show a clear pattern influencing robustness as hypothesized (see Fig. 3C and E). This is surprising as the high dropout rate was reasoned in the previous ontology-based VAE implementation as necessary when ontology terms have a large number of genes that overlap ^6,8^.

In summary, our results show the value of AUTOENCODIX, and its ontology-based or biologically-informed VAEs, for explainability of representation learning in biomedical context as well as its strength in getting methodological insights by being a platform for benchmarks.

### Towards a holistic representation: data modality translation by X-modalix

Another methodological development using VAEs is the ability of data modality translation by cross-modal autoencoders based on learning separated VAEs with aligned latent spaces for each data modality, proposed by Yang and Uhler ^5,23^. We implemented this cross-modal VAE (’X-modalix’) for two modalities including the possibility of image data integration which is unique for this AE type.

#### Efficient translation of expression data to images

To illustrate our implementation and cross-modal VAE in general, we studied two data sets and scenarios. Firstly, as a synthetic data set we used again gene expression data from TCGA pan-cancer data set and combined it with images from handwritten digits (MNIST) assigned to the five most represented cancer subtypes (see Fig. 4A). In this case, we expect the cross-modal VAE to be able to translate the gene expression signatures of cancer subtypes to the correct handwritten digits. This scenario is an illustration of the capability to translate between molecular-genetic data and image-based descriptions of cells and tissues (microscopy, radiology) which is a major goal of the field to build the bridge between genetics and phenotype.

**Figure 4:**
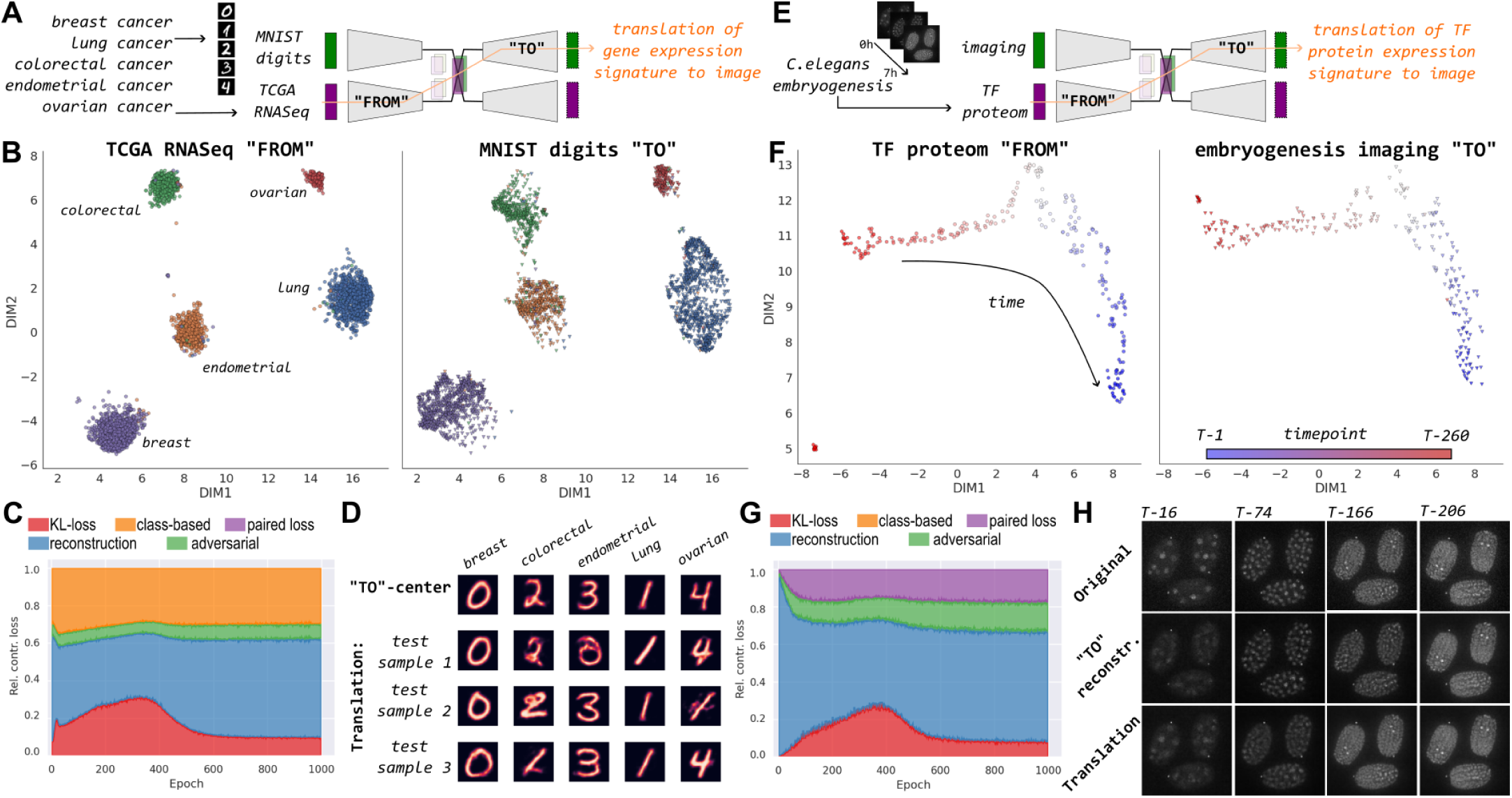
X-modalix: cross-modal VAE for translation of omics and image data for two examples: TCGA and handwritten digits (MNIST), a partly synthetic data set **(A-D)** and a real-world scenario from *C. elegans* embryogenesis **(E-H)**. 2D-UMAP representations of aligned latent spaces of each data modality for translation are shown in **B**, respectively **F**. In **C** and **G** relative contribution of loss during training is shown and calculated on a validation set using final loss term weightings (not annealed). Image translation capability is shown in **D**, respectively **H** using either the class or timepoint centers (latent dimension averages) as input for the image VAE decoder or the RNASeq (TCGA) or proteome (*C. elegans*) embedding of three randomly selected test samples, which have not been part of the training.

Secondly, we used a real-world scenario from a previously published data set of *Caenorhabditis elegans* development where livecell imaging of worms is available in combination with detailed proteome measurement of hundreds of transcription factors (see Fig. 4E). By applying our ‘X-modalix’ to this data, we expect that the transcription factor (TF) signature over 260 time points can be translated to images of *C. elegans* in the respective developmental stage.

#### Considerations and recommendations to ensure latent space alignment

For latent space alignment (learning a joint multivariate distribution) cross-modal VAEs rely on additional loss terms and training an adversarial network in parallel (see Cross-modal variational autoencoder). Our scenarios cover two cases either having non-paired data modalities (TCGA RNASeq and MNIST digits), where alignment relies on adversarial loss and semi-supervised class-based loss and paired measurements (proteome and imaging), where alignment relies on an adversarial- and a paired-loss term.

We find that in both scenarios, the correct weighting of loss terms is essential for achieving a balanced trade-off between image reconstruction capability and latent space alignment. As shown in Fig. 4C and G, we suggest defining weights (*β*, *γ*, *δ_paired_* and *δ_class_*) in a way that reconstruction loss is the majority of total loss, KL-loss less than 10% to main reconstruction precision, and the remainder controls the tightness of latent space alignment (see Fig. 4B and F).

This weighting of loss terms leads in both scenarios to well-aligned latent spaces along either classes (cancer subtypes) or time-points (developmental stages) as depicted in Fig. 4B and F. Hence, translation of gene expression signatures, respectively TF proteome data, to images works mostly very well and precise (see Fig. 4). Interestingly, even with only 260 samples (time points), the reconstruction of cellular images of *C. elegans* is very close to the original data proving the capabilities on a limited number of samples. Further, we observe only a little reduction in image reconstruction capability by latent space alignment in comparison to a purely image-based VAE (see Supplementary Figure S5).

In comparison, the class-based alignment of TCGA RNASeq data and MNIST image digits shows precise image reconstruction of class centers but bigger variations among test samples which is related to the fact that the MNIST image VAE latent space is less aligned in the outer regions of classes (see Fig. 4). We presume that this is caused by the non-paired data modalities creating a less precise alignment across and in particular within classes since there is a biological connection between data modalities in this synthetic scenario.

From this we can conclude that our implementation and cross-modal autoencoders in general, can be trained on fairly limited data sizes and both having paired measurements, but also unpaired data when having class information about samples to guide latent space alignment. It shows the immense potential of cross-modal VAEs for multi-omics studies to bridge the several layers of molecular genetic data with each other to gain insights into biological processes and diseases. Beyond, X-modalix can be used for any data-driven and multi-modal research due to its support of arbitrary images and numerical or categorical data.

## DISCUSSION

AUTOENCODIX is a versatile and generalized framework for AEs covering the full pipeline from preprocessing, training, and evaluation for common and recently emerging AE architectures. The PyTorch-based framework is open-source, easily extendable, and offers flexible hyperparameter configuration as well as in-built optimization. In connection with its pipeline characteristic covering important preprocessing steps like feature scaling and filtering, or final visualization and evaluation of embeddings, our framework can serve as an important platform for the broader application of autoencoders for representation learning not only in biomedical research but also in many other domains.

In this study, we showed the potential of our framework for testing and benchmarking AEs on pan-cancer multi-omics data as well as single-cell data. We could identify and verify important relationships like the trade-off between reconstruction capability and latent space coverage or the embedding quality in comparison to other methods like PCA or UMAP. By this comparison, we observe that disentanglement of dimension, as enforced by PCA, increases embedding quality when a larger number of dimensions are trained whereas UMAP or vanilla AEs excel in two-dimensional data representations. While we observe that ontology-based and biologically-informed VAEs support disentanglement, research studying architectures, hyperparameter influence, and loss function extension to improve disentanglement in VAEs is promising ^29–31^ and an important direction for an extension of our framework.

Further, using our in-built hyperparameter tuning via Optuna ^25^ revealed that overall improvement of reconstruction capability as well as embedding quality is very limited and requires future research for a deeper understanding and make hyperparameterization more efficient and user-friendly like other dimension reduction methods. Interestingly, we even observed for ontology-based VAEs that hyperparameter optimization may improve the training and validation loss but creates instability of the embedding towards randomness in training.

However, since this architecture should offer explainability of latent dimensions by design, this robustness is key to gaining reliable biological insights when applied to multi-omics data. The issue of robustness of biologically-informed neural networks was raised previously ^27^ and we incorporated strategies like dropout layers to counter instability by feature-ontology overlaps ^6,8^. Still, our observation that hyperparameter optimization leads to less robust embeddings suggests the need to adjust the optimization strategy to ensure reliability through repeated and randomized training. In addition, a key will be a benchmarking and deep investigation of different ontology-schemes and biological-knowledge concerning the downstream use of AE-based embeddings to maximize the quality and explainability of ontology-based VAEs ^28^. Further, besides explainability by design, post-hoc methods to determine feature importance using approaches like Shapley Additive Explanations (SHAP) ^32^, DeepLIFT ^33^ or Local Interpretable Model-agnostic Explanations (LIME) ^34^ will be a great addition to our framework to gain explainability on other AE architectures or even in combination with ontology-based VAEs.

The ability to train VAEs for the translation between data modalities has immense potential for multi-omics, but also for other domains, in a broad range of applications such as imputation of sparse and missing measurements, perturbation studies, and recovering of regulatory networks, or overall improvement representation learning of complex (biological) systems. Current implementations and use cases were tailored to specific biomedical applications, from which we generalized our cross-modal VAE and assessed its training behavior and capabilities. Based on our two scenarios, the immense potential is strengthened by its capability to translate even between very different data modalities (images and omics data), and translate even unpaired data using a semi-supervised class loss, or with very limited training data and time-series. From our experiments, it is evident that weighting the loss terms plays a vital role in training VAEs, as it facilitates proper latent space alignment while preserving the reconstruction precision of each data modality. In our opinion this warrants further research to generalize hyperparameterization of cross-modal VAEs for a broader range of applications and data modalities, in particular since loss term weights cannot be optimized via normal tuning methods using the same loss function.

In conclusion, our framework showed its value as a standardized and flexible implementation of a training and evaluation platform of AEs. Beyond its current release, it offers easy extensibility to incorporate more architectures and trends of AEs and deep-learning-based representation learning. To this end, the open-source character is key to enabling others in the community to contribute to implementing other important architectures like denoising VAE for time-series data ^19,35,36^ or to denoise count data of scRNA-seq ^37^ or for data imputation via masked VAEs ^13,38,39^. For that, we provide a contribution guide and we commit to long-term support and maintenance of the codebase for the community.

By broadening the methods and architecture range in AUTOENCODIX we would aim the application of our framework and AEs in general to other research domains as well, while on the other hand providing a platform for methodological research and benchmarking of AEs.

Finally, we envision that our framework can be used and can be a place for large-scale and pre-trained AEs and related models in biological representation learning which is the next logical step and current trend using large-scale single-cell multi-omics measurements for training ^40–43^. Such pre-trained large-scale AEs would increase the area of application in biomedical research and other domains since it either requires little sample sizes and data for fine-tuning AEs or even no training data at all for so-called zero-shot models e.g. for cell annotation and other applications.

## MATERIALS AND METHODS

### Framework and autoencoder implementation details

Our open-source framework is implemented in Python and is based on PyTorch for model implementation. It runs on Linux, Windows, and MacOS machines, preferably with GPU (CUDA) support, but also on CPU for training of AEs. Finally, we offer via Optuna ^25^ the possibility for hyperparameter tuning for crucial model and training parameters.

#### Preprocessing

Via our framework, input data is directly preprocessed to merge data modalities in accordance to AE model requirements and features are filtered and scaled upon user configuration. For the shown experiments and results, if not stated otherwise, we filtered input features (genes) for the most variance, and feature values are scaled by the in-built Scikit-Learn function of the standard scaler ^44^. We use this approach as the default matching mean squared error as reconstruction loss.

Alternatively, we offer the filtering of features also based on median absolute deviation (MAD, via SciPy package ^45^) and based on correlation using KMedoid clustering to identify representative features among highly correlated feature sets (via Scikit-Learn-Extra package ^46^, algorithm CLARA). Further, other scaling methods offered by our framework are directly embedded via Scikit-Learn such as MinMax, RobustScaler, or MaxAbsScaler).

In addition to filtering and scaling, we support categorical features through one-hot encoding and generate random train, test, and validation data splits based on user-specified ratios. For shown experiments, we used a ratio of 60:20:20 if not stated otherwise.

#### Autoencoder architectures and implementation – general

All AE implementations (see Fig. 1) are based on previously published versions and adjusted for standardization as well as generalizability and hyperparameter tuning ability. At the core of all implementations are fully-connected layers with the same series of PyTorch standard layers composed of a linear, batch normalization, drop-out (default rate: 0.1), and ReLU activation layer (if not final output layer or last layer before latent space layer). The number of layers and hidden layer dimensions in the encoder and decoder are always symmetrical and, in non-tuned variants, composed of two layers, each with a hidden layer dimension *h_i_* defined by an encoding factor *e* (non-tuned value: 4):

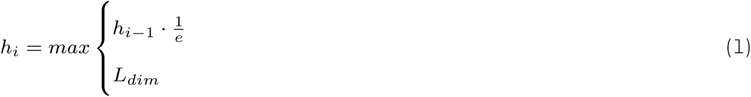

where *h_i−_*_1_ is the layer dimension of the previous layer and *L_dim_* is the user-defined dimension of the latent space.

As in previous studies and implementations of VAEs, the latent space (or the bottleneck) layer is defined by learning in parallel the mean, *µ*, and logarithmic variance, *log*(*σ*^2^), which follow a Gaussian distribution, which are combined via the reparameterization trick, such as:

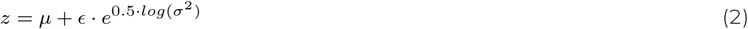

where *z* is the value of the embedding and *ɛ* a randomized value from a normal distribution. The VAE implementation, Varix, is the basis for all other VAE variants, Stackix, Ontix, and X-Modalix.

Training of all AE architectures is based on the minimization of the reconstruction loss, with additional loss terms depending on the architecture and described in the following. For the results shown, we employed the mean squared error (MSE) as the reconstruction loss, since it matches our input data characteristics. However, as an alternative, we implemented the option to use binary cross entropy (BCE) for reconstruction loss as well (both via in-built PyTorch functions). For all VAEs similarity loss to normal distribution is determined by the Kullback–Leibler (KL) divergence as default and alternatively, users can opt for Maximum Mean Discrepancy (MMD), which are both weighted via the hyperparameter *β* to control trade-off to the reconstruction loss.

All implementations are open-source with a special focus on standardization and modularization to enable flexible and easy extension of the framework to other emerging AE architectures and variants.

#### Stacked variational autoencoder

The stacked VAE (Stackix), also referred to as a hierarchical VAE, is has the core idea on which we built our implementation that separate VAEs are first trained on each of the data modalities before being merged together ^22^. The resulting latent space embeddings of each VAE have a dense layer of dimension 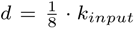,*m*, where *k_input,m_* is the number of input features for data modality *m*. These dense layer embeddings are the input for the stacked VAE with the final latent dimension *L_dim_* as specified by the user.

#### Ontology-based variational autoencoder

The ontology-based VAE, (Ontix), aims to incorporate biological or other domain knowledge into the architecture. We follow current ideas of knowledge incorporation ^6,8^ by making the decoder layers sparse and mask according to the feature-ontology term or ontology-ontology term connectivity. In our implementation, as in previous, we enforce zero weights in the respective linear layers to mask and restrict the decoder. Current implementations of ontology-based VAE are either single (linear) layer decoders (VEGA ^6^, expiMap ^7^) or can have any depth of ontology (OntoVAE ^8^). For a trade-off of simplicity and non-linearity complexity, our implementation supports either one or two levels of feature-ontology hierarchies. In addition, users can specify additional, fully-connected layers if additional dimension reduction (e.g. 2D visual representation) is necessary.

#### Cross-modal variational autoencoder

The X-Modalix implementation consists of two parts: (a) the method of training two autoencoders together to enable the translation functionality and (b) the model architecture itself. For (a), we followed closely the implementation of Yang et al. ^23^ and made adjustments for standardization and generalization to match our framework approach. The general idea of a cross-modal VAE is to translate data modality *A* to data modality *B* by putting *A* in the encoder part of the VAE for modality *A*, obtaining the latent representation *z*, and feeding *z* into the decoder of the VAE for data modality *B*. In our case, we always translated numeric omics data into image data. To ensure a functional translation process, the different autoencoders need to be trained together to keep the latent spaces aligned.

For (b), we used the Varix architecture (as described above in Section Autoencoder architectures and implementation - general) for numerical data and combined it as described in (a) with a specialized VAE for image data. The image VAE processes an input image of shape (*C, H, W*), where *C* is the number of channels, and *H* = *W* (we allow only quadratic images as of now) represents the height and width of the image, respectively, and reconstructs the same-sized output image. The architecture consists of an encoder and a decoder of convolutional layers, connected via a latent space representation.

In line with previous image VAE implementations ^23^, the encoder consists of five convolutional layers, each with a kernel size of 4, stride of 2, and padding of 1.

We offer pretraining of the image VAE to account for the different complexities of the data modalities, as described by Yang *et al.* ^23^, for a user-specified number of epochs before the combined training of both VAEs with latent space alignment. Further, we experienced that pretraining of the image VAE combined with *β* annealing increases robustness towards posterior collapse during latent space alignment. For latent space alignment, the loss function consists of five different terms (i-v): the reconstruction loss (i) and similarity loss (ii) identically to all VAEs. In addition, an adversarial loss term (iii), weighted with a parameter *γ*, as well as a paired loss term, weighting *δ_paired_* (iv) and a semi-supervised class-based loss term, weighting *δ_class_* (v). The combination of (iii)-(v) is crucial for latent space alignment and subsequent data modality translation.

To obtain the adversarial loss term, a latent space classifier is trained in parallel that learns to classify whether the latent space comes from the image- or omics data modality. We used cross entropy as a loss function for this term. In the loss term for the VAEs, we then switch the labels of the data modalities in the cross-entropy function. This results in a higher loss when the latent spaces are easy to discriminate for the latent space classifier and thus keeps the latent spaces aligned. The latent space classifier itself is a fully-connected neural network consisting of an input layer, two hidden layers, and an output layer.

The paired loss term, or the semi-supervised class-based loss term, improves latent space alignment and compactness by minimizing the distance, defined as the mean absolute deviation across all latent dimensions, of a sample across both latent spaces per data modality (paired) or samples to its class average (center, e.g. cancer subtype). Since paired measurements of data modalities are rare in multi-omics studies, the class-based approach is an important alternative for real-world applications.

### Data sets

To test and benchmark the capabilities of our framework we use the following publicly available data sets to mimic typical use cases and provide a reference for use on other data sets and applications.

(*TCGA*) Pan cancer multi-omics data set: The Cancer Genome Atlas (TCGA) ^47^, retrieved as preprocessed files via cBioPortal ^48^, include patient data for a total of 32 cancer types. We use three data modalities: gene expression data (mRNA-seq V2 RSEM), epigenetic data (methylation HM27 and HM450), and mutational data by combining single-nucleotide variations and copy-number alterations by a simple score, *M_score,i_*, per gene *i*:

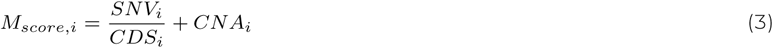

where *SNV_i_* is the number of single-nucleotide variations per gene, *CDS_i_* the length of the gene (coding region) and *CNA_i_* the copy number alteration of the gene. In total, the data set contains 9267 patient samples with data files across all three modalities.

(*sc-Cortex*) Single-cell multi-omic (paired RNA-Seq and ATAC-Seq) data set of the developing human cerebral cortex ^49^: retrieved as preprocessed h5ad files from CZ CELLxGENE Discover ^50^. Includes a total of 45,549 cells (nuclei).

*C. elegans* embryogenesis microscopic images and proteomic data, based on transcription factor reporter atlas, were used for the cross-modal VAE, ^51^. Further, MNIST handwritten digit images retrieved via the Keras package ^52^ were used in combination with the TCGA gene expression data.

A summary of data sets and key figures is given in the Supplementary Table S1.

### Experiments

All of the scripts for the shown experiments and the visualizations of their results can be accessed and reproduced via our git repository, https://github.com/jan-forest/autoencodix-reproducibility. In particular, the benefit of our pipeline is the configuration via YAML-files providing reproducibility by design as well as usage on new data sets.

#### Impact of *β*-annealing and training of variational autoencoders

As a baseline representation, a Varix with two-dimensional latent space, which enables direct visualization, was trained on all three data modalities of the TCGA data set with 2000 features (genes) of each modality. During training for over 2000 epochs, weighting factor *β* of the KL-divergence loss term was altered every 400 epochs to analyze the impact of KL loss term on training and latent space of VAEs. We implemented the option to control *β*-annealing via different functions, and, for the shown experiment, we use a function with five constant phases:

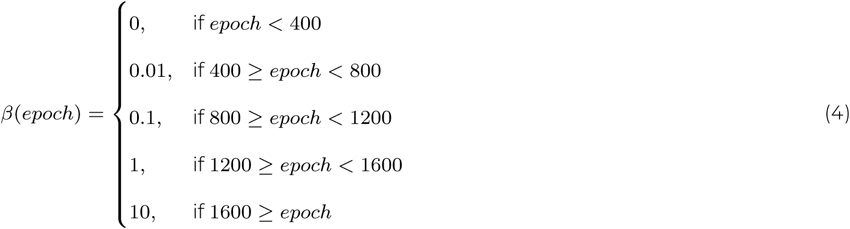

If not stated otherwise, the default *β*-annealing function with a simple logistic annealing is used:

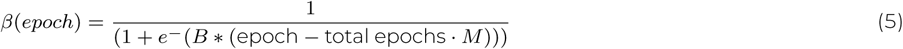

and provides robustness of VAEs training towards posterior collapse as shown previously ^53,54^.

When using VAEs as generative models, a good latent space coverage would be critical to generate synthetic and meaningful samples in a broad range. To define the latent space coverage, we assume the following, given the samples are perfectly distributed among latent space dimensions; if the latent space is only one-dimensional, one could have a grid with *b* bins where *b* equals the number of samples (*n*) and when perfectly distributed each bin would contain exactly one sample. If the latent space is two-dimensional, each latent dimension can be binned with 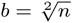 or, in general, 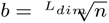 for any latent dimension. Bins to check coverage are then defined as a grid between minimal and maximal values for each latent dimension across samples. The coverage of a latent dimension *l_i_* is then defined as the fraction of bins containing at least one sample to the total number of bins *b*. In Fig. 2B the average coverage over the two dimensions is shown. For Gaussian distributions, a maximal coverage of 0.5 can be expected.

#### Comparison of autoencoder architectures and the impact of tuning for multi-omics data integration

For benchmarking of the available AE architectures of our framework, we used the TCGA and sc-Cortex data set and trained AE on all combinations of up to three data modalities with a uniform configuration and parameters, as outlined in table S2. Furthermore, key hyperparameters (see table S2) have been tuned via Optuna as an in-built option of our framework for the same number of epochs and 50 optimization trials. Data sets were split into training, validation, and test sets in the ratio of 60:20:20.

The ontology-based VAE (Ontix) was constructed and trained using biological pathways from the Reactome database ^26^. The highest level of pathway hierarchy (top-level) was used to define the 29 dimensions in the latent space. As for the hidden-layer, we used the third-highest level in the pathway hierarchy to define the first ontology level between the genes and each pathway. When compared to other AE architectures with *L_dim_* = 2 or 8, a third hidden-layer (fully-connected) is introduced for dimensionality reduction and comparability to other architectures with *L_dim_ <* 29.

To evaluate embedding (AEs, PCA or UMAP) performance, available biological and clinical annotation of samples were used to train regression or classification tasks using three different machine learning algorithm classes (linear or logistic regression, support vector machines with radial kernel, and random forests) using scikit-learn implementations. The following performance metrics were used: ROC AUC for classification tasks and explained variance *r*^2^ for regression tasks. To enable comparability across tasks with varying difficulty, the performance using randomly selected features as the simplest mean of dimension reduction (five repetitions) was determined.

The performance of embeddings over randomly selected features (*z_i,m_*) is then calculated and determined based on Z-score normalization to account for the mean (*avg*) and standard deviation (*sd*) across five (*r* ∈ [1, 5]) randomly selected features *η_r,i,m_* for the *i*-th classification or regression task and *m*-th machine learning algorithm:

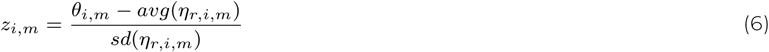

where *θ_i,m_* is the performance of the embedding (autoencoder, PCA or UMAP) as either explained variance (*r*^2^, regression tasks) or ROC AUC (classification tasks). We apply this Z-score normalization in order to take into account that some machine learning tasks, such as cell type classification, are easier and expected improvements from embeddings are much smaller than for hard prognostic tasks, such as months of survival.

#### Analysis and robustness of ontology-based autoencoders

To test the reliability and robustness of trained latent space intensities associated with ontologies, we conducted five repetitions per hyperparameter and ontology combination. As ontologies, we used Reactome pathways (as described in previous sections) and chromosomal positions where latent space dimensions represent human chromosomes (including X,Y, and mitochondrial genes). As a hidden layer, we used information via Ensembl BioMart ^55^ about cytobands to infer substructures of human chromosomes (version GRCh38.p14) where genes (features) are located.

The baseline hyperparameter configuration is *β* = 0.1, *p_dropout_* = 0.5, and a of learning rate 1*e* − 4. To gain insights into the impact of these parameters on robustness, we varied each parameter individually for five different values and performed five training repetitions. Each training is randomized across the whole pipeline including train, test, and validation splits or model weight initialization.

As a measurement of robustness, we calculate the average Pearson correlation across all repetitions for each latent space dimension representing an ontology. Since we are not interested in qualitative robustness and have not reversed orders in latent spaces, we calculate the absolute value of correlation before averaging over repetition combinations.

#### Scenarios of cross-modal autoencoder

To showcase the capabilities of the X-modalix, we ran two experiments: First, we investigate the cross-modal translation from expression data of transcription data to embryonic images of the *C. elegans* organism. Second, we synthetically coupled TCGA omics data with handwritten digits (MNIST).

The first scenario uses a protein expression atlas of transcription factors (TFs) at single-cell resolution, collected by Ma et al. ^56^, mapped onto developmental cell lineages during *C. elegans* embryogenesis as well as corresponding 4D imaging of embryogenesis. The authors took images every 75 seconds during the embryonic development. At each time point, three slide positions were scanned across 30 focal (z) planes with a spacing of 1 µm between planes. This results in 300 images (one for each time point) per TF. Each image has 30 z-channels and two color channels, which we processed as follows:

We used, for simplicity, only one channel (red) and applied maximum intensity projection across the z-dimension resulting in 512×512 images per timepoint and TF. Pixel values are normalized to a range of 0 to 255 and TF gene ALY-2 was selected based on its strong signal across all timepoints and cell states.

Proteomic expression of TF is aggregated and normalized across all strains and time points. We normalized the expression value of each TF by the sum of the expression of all TF at the same time point to retrieve a proteomic signature rather than absolute expression across time points. After removing missing values (dropping TFs consisting of more than 90% missing values and then removing all time points with missing values) the dataset had 260 time points (samples) and 290 TFs (features).

We used a 70% train, 10% validation, and 20% test split for the dataset. The image VAE was pretrained for 100 epochs including logistic *β*-annealing. We show a complete list of hyperparameters in the Supplementary Table S2. We optimized the loss term weights *β*, *γ*, *δ_paired_*, and *δ_class_* manually to achieve a balance between reconstruction capability and latent space alignment.

For the second cross-modal experiment, we mapped MNIST handwritten digit images to five most common TCGA cancer types. We processed the TCGA dataset as before using gene expression data modality only. We obtained the MNIST dataset from the Keras package ^52^ and mapped the digits zero to four to the most frequent cancer types ("Breast Cancer": 0, "Non-Small Cell Lung Cancer": 1, "Colorectal Cancer": 2, "Endometrial Cancer": 3, "Ovarian Epithelial Tumor": 4,). The resulting dataset consists of 3230 images, with 978 images for the digit zero, 966 images for one, 522 for two, 563 images for three, and lastly, 201 for the digit four. Hyperparameters and configuration is summarized in Supplementary Table S2. We evaluated the translation as in the *C. elegans* example. Additionally, we defined centers of cancer subtype classes by calculating the latent space embeddings of all samples of each class for either the RNASeq VAE encoder ("FROM") or the image-based VAE ("TO"). When retrieving center representations from image-based VAE decoder ("TO"), for each class and each latent dimension, the median embedding value across samples was taken as input for the decoder.

A quantitative evaluation of the image reconstruction quality is given in Supplementary Figure S5 for both scenarios.

## ACKNOWLEDGMENTS

MJ acknowledges support by the German Federal Ministry of Education and Research under the funding code 03ZU1111MB (SaxoCellOmics) as part of the Clusters4Future cluster SaxoCell. JE and NJ acknowledge financial support by the Federal Ministry of Education and Research of Germany and by Sächsische Staatsministerium für Wissenschaft, Kultur und Tourismus in the programme Center of Excellence for AI-research „Center for Scalable Data Analytics and Artificial Intelligence Dresden/Leipzig“, project identification number: ScaDS.AI. DP acknowledges support by the Sächsische Staatsministerium für Wissenschaft, Kultur und Tourismus (SMWK) under the frame of ERA PerMed (GRAMMY, 2019-275 and MIRACLE, 2021-055).

Further, we thank Isabella Kreller and all colleagues from ScaDS.AI Dresden/Leipzig for framework testing and critical feedback. NJ recognises the editing guidance from the Kather group, EKFZ, TU Dresden.

The TCGA data set used in our experiments was generated by the TCGA Research Network: https://www.cancer.gov/tcga.

## AUTHOR CONTRIBUTIONS

MJ and JE mainly developed the software, conceived the study design and data analysis. NJ and DP supported in development of the software and data analysis. NS supported in the study design and framework conceptualization. MJ and JE wrote the manuscript draft and all authors contributed in writing the final manuscript including visualizations and repository preparation.

## SUPPLEMENTARY INFORMATION

### Data Sets

**Table S1:** Overview of data sets used in experiments.

| Data set name | Reference | Data modality | # samples | # features | pre-filtering | subtypes |
| --- | --- | --- | --- | --- | --- | --- |
| TCGA |  | RNASeq | 10059 |  | no | 32 cancer subtypes |
|  |  | Mutational score | 9866 |  | no |  |
|  |  | DNA Methylation | 10729 |  | no |  |
|  |  | combined | 9267 |  |  |  |
| sc-Cortex |  | sc-RNASeq | 24437 | 6000 | yes, via scanpy | 15 cell types |
|  |  | sc-ATACSeq | 45549 | 6000 | yes, via scanpy |  |
|  |  | combined | 24437 |  |  |  |
| Celegans |  | Proteome | 260 | 279 | no | 260 timepoints |
|  |  | Image | 260 | 128x128 pixel | no | 260 timepoints |
| MNIST |  | Image | 3230 | 64 x 64 pixel | no | 0-4 digits |

### Overview experimental set-up and hyperparameterization

**Table S2:**
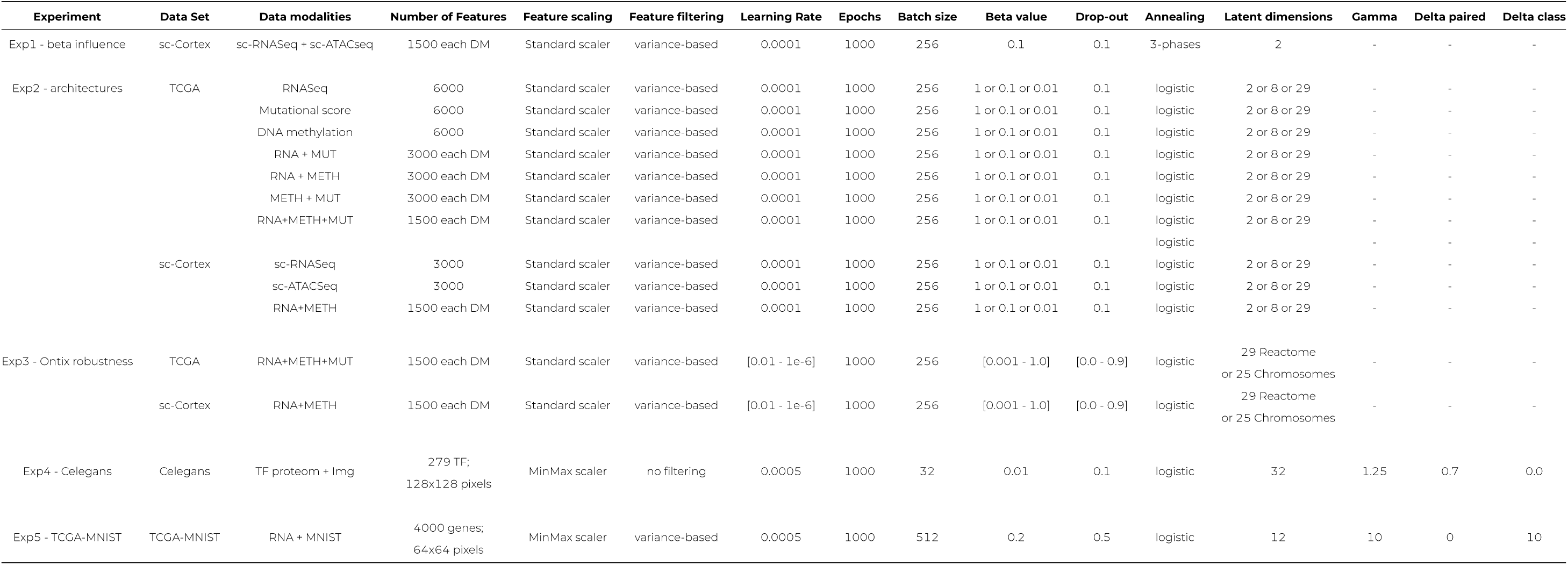
Overview of experiments and hyperparameter configuration.

### Details architecture comparison

**Figure S1:**
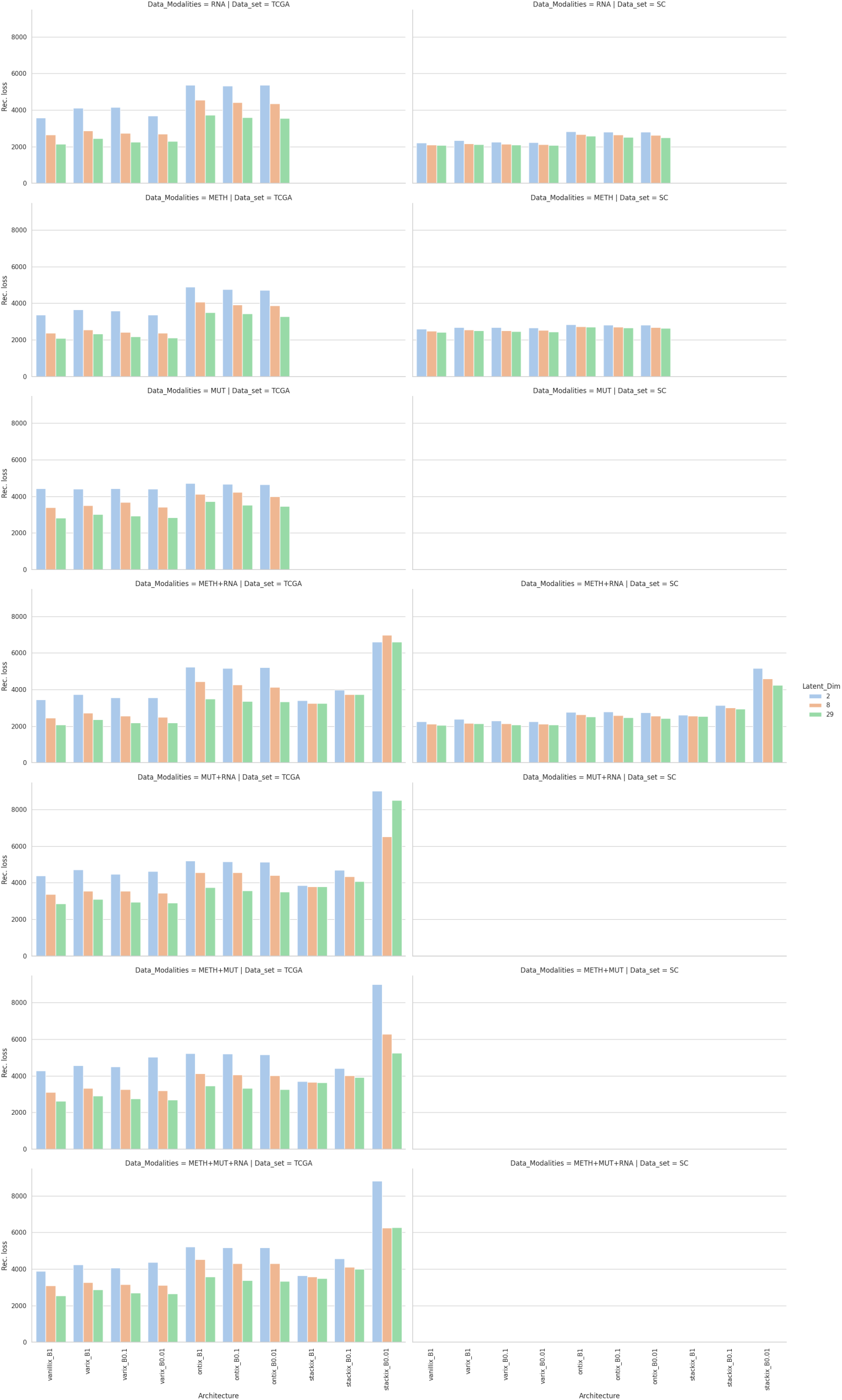
Detailed reconstruction capability of AE types

**Table S3:**
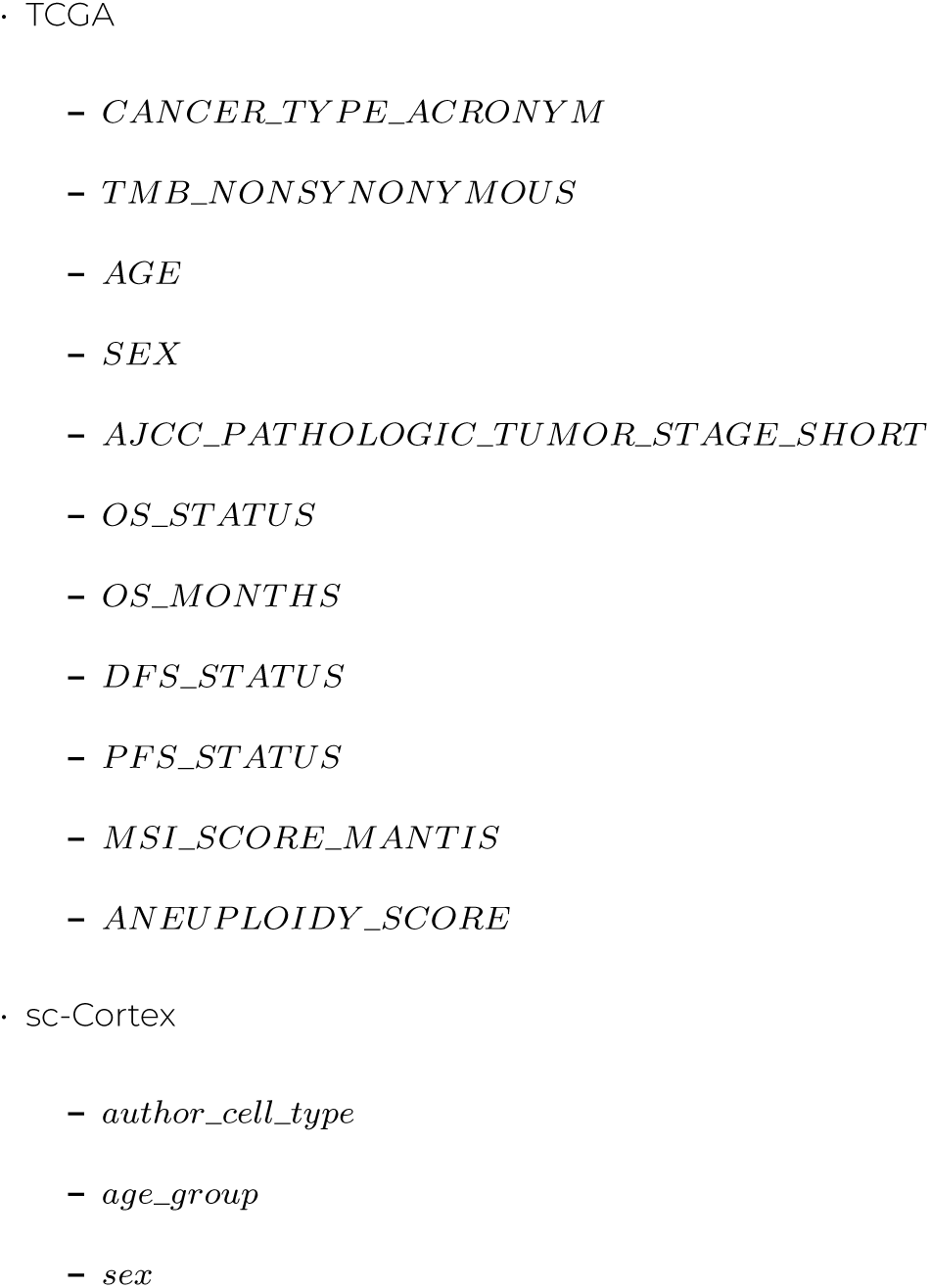
Overview of tasks (annotation labels) for supervised learning on embeddings.

**Figure S2:**
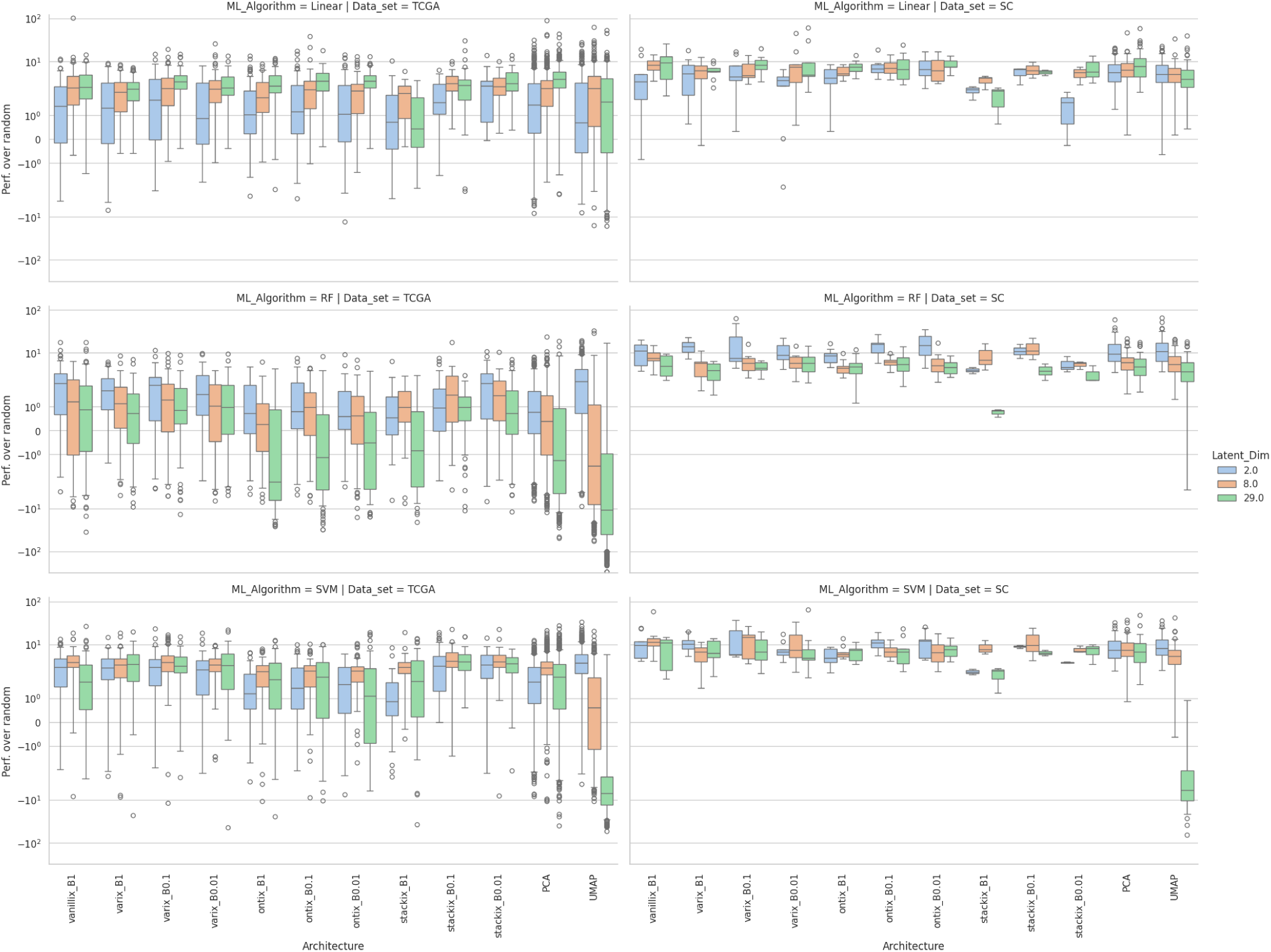
Detailed embedding performance of AE types

**Figure S3:**
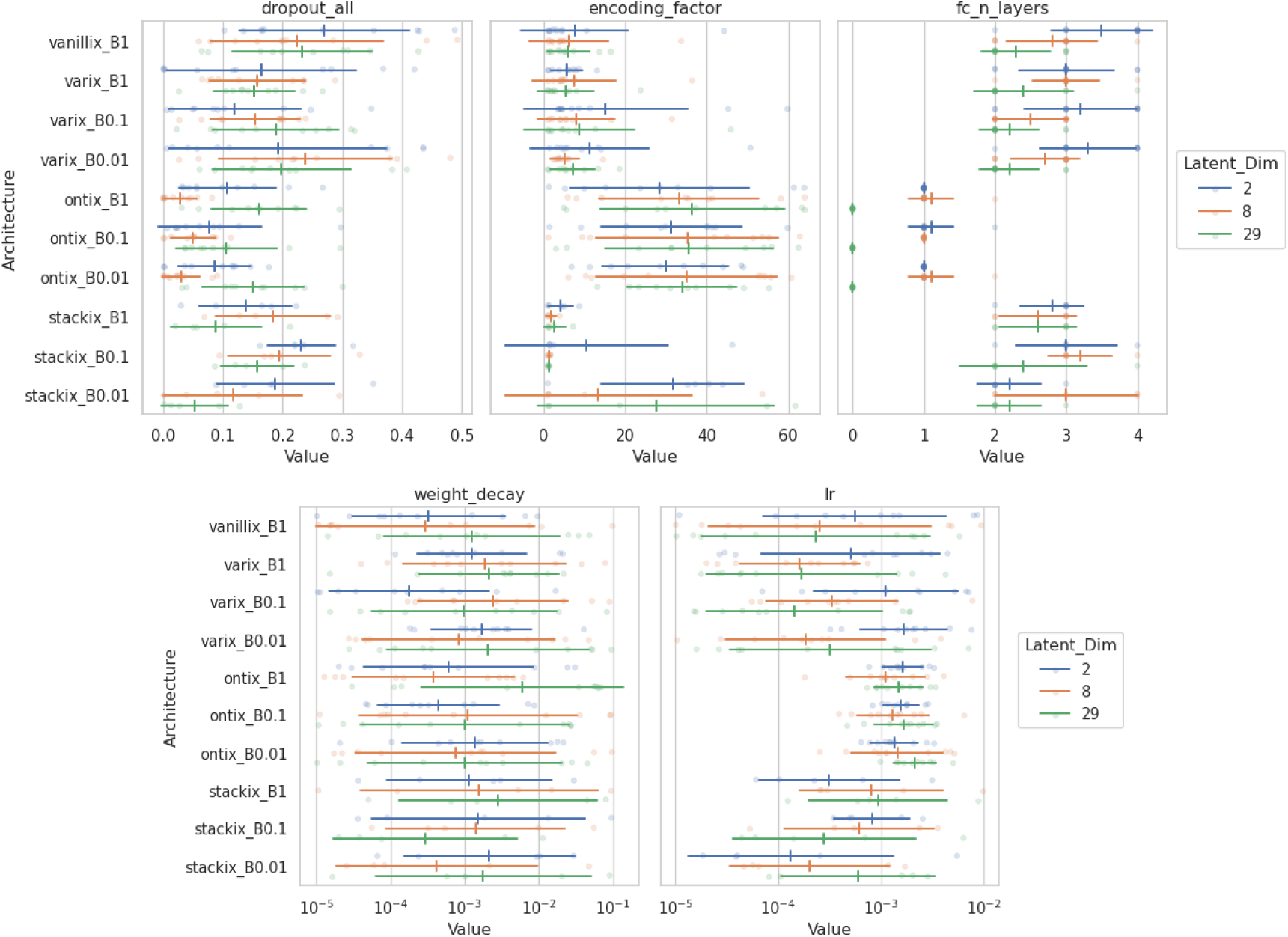
Value range of optimized hyperparameters after tuning.

### Ontology-based VAE on single-cell data

**Figure S4:**
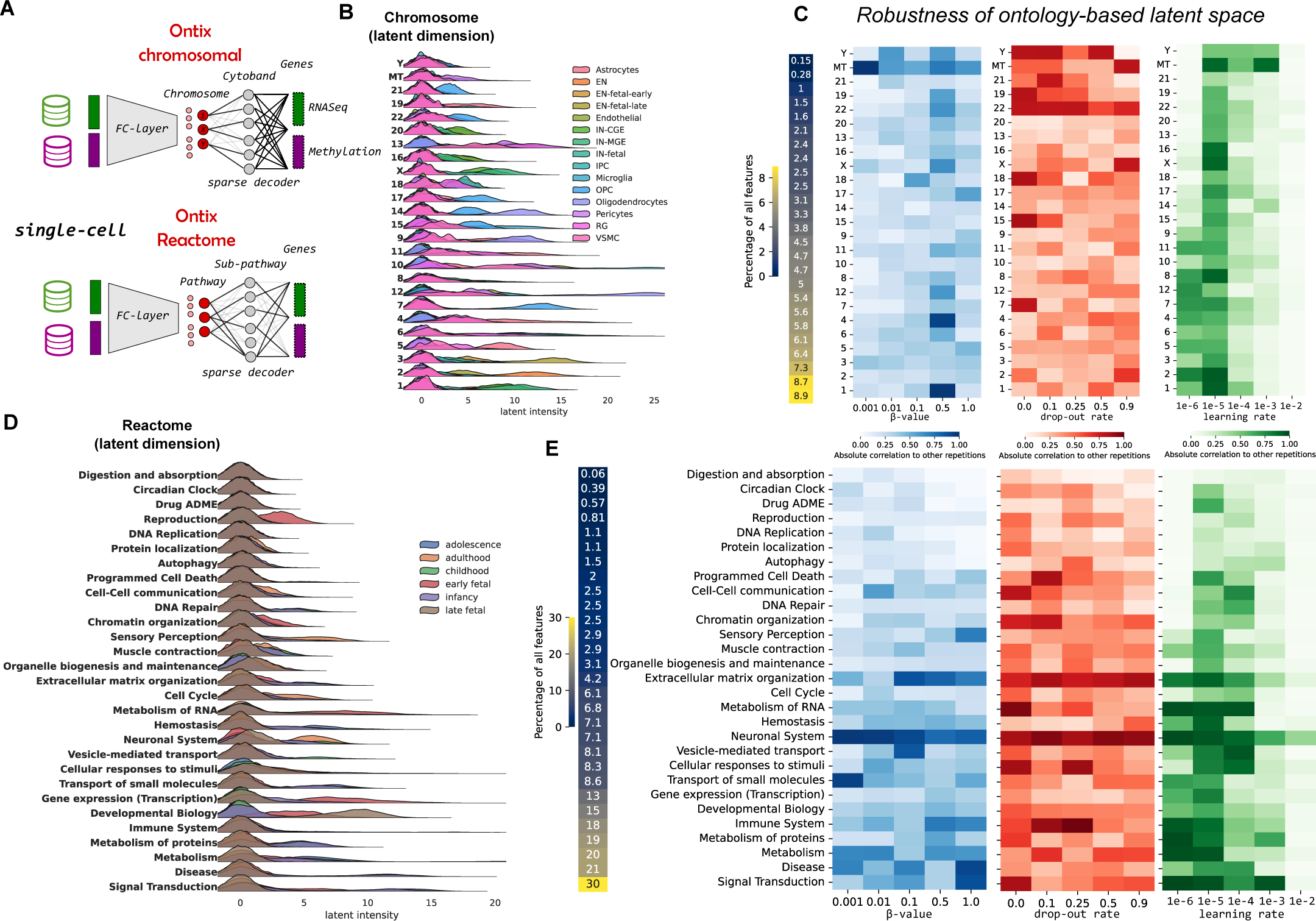
Ontix: biological-informed decoder enables explainability of latent space, but its robustness depends on hyperparameters. **(A)** Two types of ontologies are tested on single-cell data of cortical development: chromosomal-based and Reactome-pathway-based latent dimensions. **(B)** and **(D)** show ridge-line plots of latent intensity distributions based for selected examples of sample classes (cell type and age developmental groups) to show applicability for explainability. **(C)** and **(E)** show the robustness of these embeddings in dependence of hyperparameters. Robustness is defined as the mean absolute Pearson correlation between five independent training runs with randomized data split and weight initialization.

### Cross-modal VAE evaluation details

To quantify the translation capabilities of the cross-modale autoencoder we implemented two evaluation experiments. Firstly, we compared the reconstructions of three different cross-modal autoencoders with the original images by calculating the pixel-wise MSE. The three architectures were: (a) the actual cross-modal autoencoder as explained in the Material and Methods sections, (b) a "fake" cross-modal autoencoder that translates images to images (the training process was the same as for the true cross-modal autoencoder with the sole difference that both data modalities were the *C. elegans* images — we refer to this experiment as img_img_pure), and (c) the actual cross-modal autoencoder, but we feed the test images into the network (in contrast to feeding the test proteomics data and obtaining images). Figure S5 **A** shows the aggregated MSEs over all time points as a boxplot. Figure S5 **B** shows the MSE for each timepoint reconstruction for the three different experimental setups. The MSE for the "true" cross-modal autoencoder is similar to the two compare-group-autoencoders. This indicates that the translation capabilities are potent.

**Figure S5:**
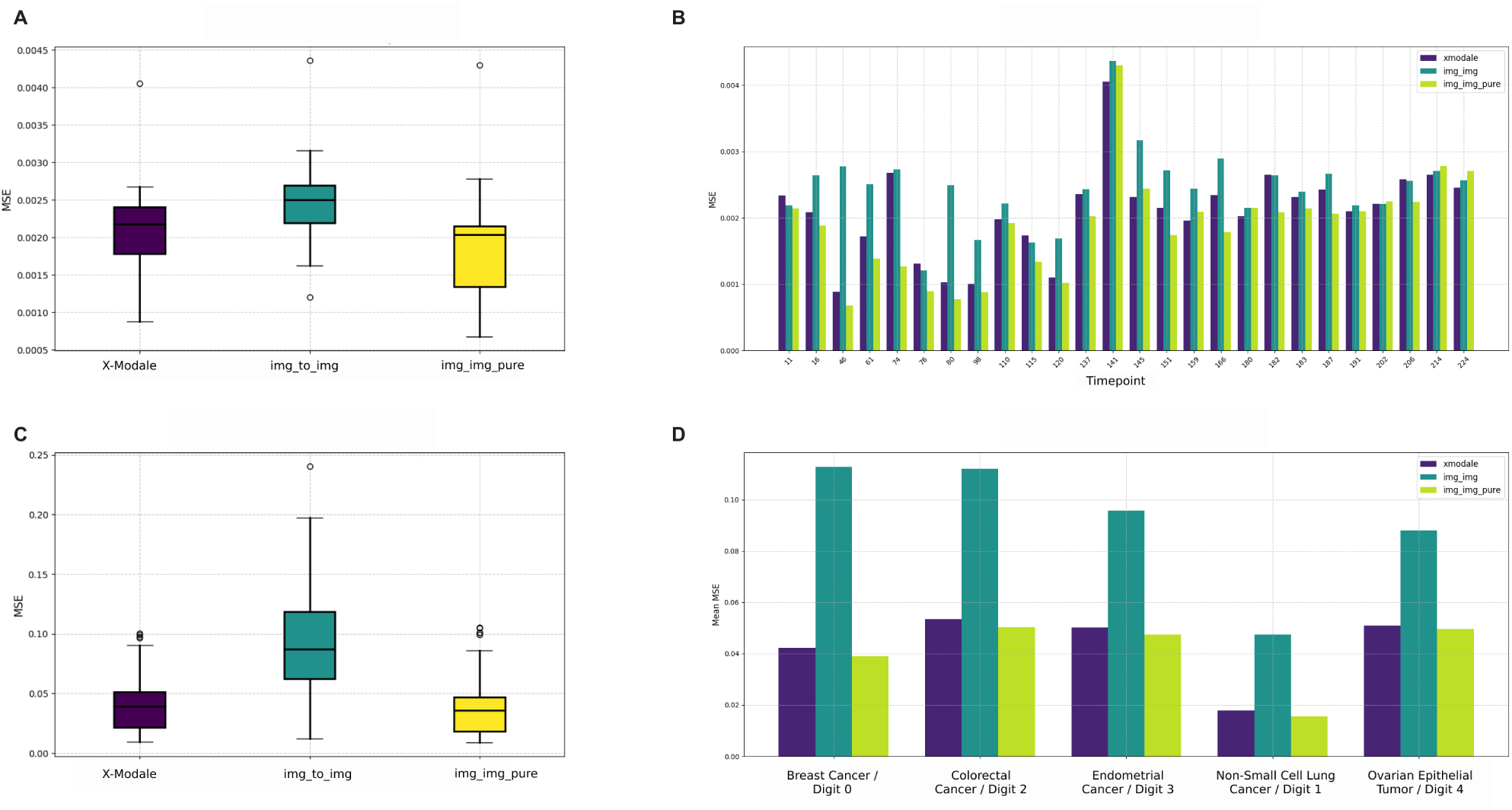
Quantitative evaluation of the cross-modal autoencoder. **(A)** MSE of all predictions of the test samples compared with the original images for different autoencoder scenarios depicted as boxplot for the C*C. elegans* experiment. i**(B)** MSE between original and reconstructed test samples at each timepoint for three different autoencoder scenarios for *C. elegans* experiment.. **(C)**See figure A, but for the TCGA/MNIST example **(D)**See figure B, but for the TCGA/MNIST example.

Secondly, we trained a basic convolutional neural network (CNN) regression model on the original training images to predict the time point of each image. The rationale behind this approach was that if the translation performs effectively, the trained model should achieve similar performance when inferring time points from both the original test data and the reconstructed test data. Model performance was evaluated using the R² metric, as reported in Table S4. Additionally, we also trained the CNN with the reconstructed train data and evaluated it with the reconstructed test data, to investigate whether the differences in performance come from a distribution shift, or from changed task difficulty (shown in the row Test Reconstructed (trained_recons))

**Table S4:** Model evaluation R2 metric across datasets.

| Dataset | $R^2$ |
| --- | --- |
| Train Original | 0.9460 |
| Train Reconstructed | -0.9351 |
| Validation Original | 0.9140 |
| Validation Reconstructed | -0.9355 |
| Test Original | 0.9200 |
| Test Reconstructed | -1.6367 |
| Test Reconstructed (trained_recons) | 0.2564 |

The architecture of the CNN model used for regression consists of three convolutional layers followed by fully connected layers. Each convolutional layer applies filters to extract features from the input images. Specifically, the first layer employs 16 filters of size 3×3, followed by a ReLU activation function and a 2×2 max-pooling operation. This is repeated for subsequent layers, which increase the number of filters to 32 and then 64. The output from the final convolutional layer is flattened into a 1D vector of size 64×28×28 and passed through two fully connected layers: the first with 128 units and a ReLU activation, and the second layer producing a single scalar output for regression. The model was trained for 50 epochs using mean squared error (MSE) as the loss function.

The results presented in Table S4 offer a clear comparison of the model’s performance on the original and reconstructed datasets. For both the training and test sets, the *R*^2^ values for the original images were consistently high (*R*^2^ *>* 0.9), indicating that the model effectively learned to predict the time points from the original images. However, the performance on the reconstructed data was significantly poorer, with negative *R*^2^ values. This disparity suggests that the reconstructed images originate from a different distribution than the original images, leading to the model’s inability to generalize effectively. When considered alongside the results depicted in Figures S5 **A** and **B**, we conclude that while the cross-modal translation method performs comparably to traditional reconstruction techniques, it still alters the data distribution in a way that impairs the ability of AI models to learn accurately.

We performed the same experiments for the TCGA/MNIST example as shown in Figures S5 **C** and **D** and Table S5. The only difference is that we trained a classifier instead of a regression model. The architecture for this CNN classifier closely mirrors that of the regression model, featuring the same three convolutional layers for feature extraction. However, the final fully connected layer was adjusted to output probabilities for four discrete classes, representing different cancer types. As assessed by the F1 score (see Table S5), the model’s performance was significantly better on the original datasets than the reconstructed ones.

Table S5 shows that the classifier achieved an F1 score of 0.9837 on the original training data and 0.9754 on the original test data, indicating high classification accuracy. However, on the reconstructed datasets, the F1 score dropped to 0.7031 for the training set and 0.6791 for the test set, showing a similar trend to the regression experiments — but not as stark. This is likely because classifying digits is easier than inferring time points.

**Table S5:** F1 Score of models on various datasets.

| Dataset | F1 Score |
| --- | --- |
| Train Original | 0.9837 |
| Train Reconstructed | 0.7031 |
| Validation Original | 0.9667 |
| Validation Reconstructed | 0.7067 |
| Test Original | 0.9754 |
| Test Reconstructed | 0.6791 |
| Test Reconstructed Trained on Reconstructed | 0.7161 |

A noteworthy observation from both experiments is that even when the model was trained on the reconstructed images, the regression and classification performance still decreased compared to the original data. This could indicate that the reconstructions do not preserve enough detail and accuracy, which negatively impacts the model’s ability to learn and generalize effectively.

#### Class Frequencies

The datasets contain different numbers of cases across four cancer types. In the training set, there are 648 Non-Small Cell Lung Cancer cases, 706 Breast Cancer cases, 396 Endometrial Cancer cases, 367 Colorectal Cancer cases, and 144 Ovarian Epithelial Tumor cases. In the validation set, there are 223 Non-Small Cell Lung Cancer cases, 181 Breast Cancer cases, 107 Endometrial Cancer cases, 95 Colorectal Cancer cases, and 40 Ovarian Epithelial Tumor cases. For the test set, there are 95 Non-Small Cell Lung Cancer cases, 91 Breast Cancer cases, 60 Endometrial Cancer cases, 60 Colorectal Cancer cases, and 17 Ovarian Epithelial Tumor cases.

## REFERENCES

[1] Milan Picard, Marie-Pier Scott-Boyer, Antoine Bodein, Olivier Périn, and Arnaud Droit. Integration strategies of multi-omics data for machine learning analysis. Computational and Structural Biotechnology Journal, 19:3735–3746, 2021.

[2] Mingon Kang, Euiseong Ko, and Tesfaye B Mersha. A roadmap for multi-omics data integration using deep learning. Briefings in Bioinformatics, 23(1):bbab454, 2022.

[3] Pengzhi Li, Yan Pei, and Jianqiang Li. A comprehensive survey on design and application of autoencoder in deep learning. Applied Soft Computing, 138:110176, 2023.

[4] Kamal Berahmand, Fatemeh Daneshfar, Elaheh Sadat Salehi, Yuefeng Li, and Yue Xu. Autoencoders and their applications in machine learning: a survey. Artificial Intelligence Review, 57(2):28, 2024.

[5] Karren D Yang and Caroline Uhler. Multi-domain translation by learning uncoupled autoencoders. arXiv preprint arXiv:1902.03515, 2019.

[6] Lucas Seninge, Ioannis Anastopoulos, Hongxu Ding, and Joshua Stuart. Vega is an interpretable generative model for inferring biological network activity in single-cell transcriptomics. Nature Communications, 12(1):5684, 2021.

[7] Mohammad Lotfollahi, Sergei Rybakov, Karin Hrovatin, Soroor Hediyeh-Zadeh, Carlos Talavera-López, Alexander V Misharin, and Fabian J Theis. Biologically informed deep learning to query gene programs in single-cell atlases. Nature Cell Biology, 25(2):337–350, 2023.

[8] Daria Doncevic and Carl Herrmann. Biologically informed variational autoencoders allow predictive modeling of genetic and drug-induced perturbations. Bioinformatics, 39(6):btad387, 2023.

[9] Tianle Ma and Aidong Zhang. Integrate multi-omics data with biological interaction networks using multi-view factorization autoencoder (mae). BMC Genomics, 20(Suppl 11):944, 2019.

[10] Muta Tah Hira, MA Razzaque, Claudio Angione, James Scrivens, Saladin Sawan, and Mosharraf Sarker. Integrated multi-omics analysis of ovarian cancer using variational autoencoders. Scientific Reports, 11(1):6265, 2021.

[11] Edian F Franco, Pratip Rana, Aline Cruz, Victor V Calderon, Vasco Azevedo, Rommel TJ Ramos, and Preetam Ghosh. Performance comparison of deep learning autoencoders for cancer subtype detection using multi-omics data. Cancers, 13(9):2013, 2021.

[12] Richard Lupat, Rashindrie Perera, Sherene Loi, and Jason Li. Moanna: multi-omics autoencoder-based neural network algorithm for predicting breast cancer subtypes. IEEE Access, 11:10912–10924, 2023.

[13] Lei Huang, Meng Song, Hui Shen, Huixiao Hong, Ping Gong, Hong-Wen Deng, and Chaoyang Zhang. Deep learning methods for omics data imputation. Biology, 12(10):1313, 2023.

[14] Mengke Guo, Xiucai Ye, Dong Huang, and Tetsuya Sakurai. Robust feature learning using contractive autoencoders for multi-omics clustering in cancer subtyping. Methods, 2024.

[15] Pedro Henrique da Costa Avelar, Le Ou-Yang, Min Wu, and Sophia Tsoka. Pathway activity autoencoders for enhanced omics analysis and clinical interpretability. In IEEE International Conference on Bioinformatics and Biomedicine, 2024.

[16] Yehudit Hasin, Marcus Seldin, and Aldons Lusis. Multi-omics approaches to disease. Genome Biology, 18:1–15, 2017.

[17] John N Weinstein, Eric A Collisson, Gordon B Mills, Kenna R Shaw, Brad A Ozenberger, Kyle Ellrott, Ilya Shmulevich, Chris Sander, and Joshua M Stuart. The cancer genome atlas pan-cancer analysis project. Nature Genetics, 45(10):1113–1120, 2013.

[18] Debabrata Acharya and Anirban Mukhopadhyay. A comprehensive review of machine learning techniques for multi-omics data integration: challenges and applications in precision oncology. Briefings in Functional Genomics, page elae013, 2024.

[19] N Costa, L Pérez, and L Sánchez. Rapidae: A python library for creation, experimentation, and benchmarking of autoencoder models. In 2024 IEEE International Conference on Fuzzy Systems (FUZZ-IEEE), pages 1–8. IEEE, 2024.

[20] Adam Paszke, Sam Gross, Francisco Massa, Adam Lerer, James Bradbury, Gregory Chanan, Trevor Killeen, Zeming Lin, Natalia Gimelshein, Luca Antiga, et al. Pytorch: An imperative style, high-performance deep learning library. Advances in neural information processing systems, 32, 2019.

[21] Isaac Virshup, Sergei Rybakov, Fabian J Theis, Philipp Angerer, and F Alexander Wolf. anndata: Annotated data. bioRxiv, pages 2021–12, 2021.

[22] Nikola Simidjievski, Cristian Bodnar, Ifrah Tariq, Paul Scherer, Helena Andres Terre, Zohreh Shams, Mateja Jamnik, and Pietro Liò. Variational autoencoders for cancer data integration: design principles and computational practice. Frontiers in Genetics, 10:1205, 2019.

[23] Karren Dai Yang, Anastasiya Belyaeva, Saradha Venkatachalapathy, Karthik Damodaran, Abigail Katcoff, Adityanarayanan Radhakrishnan, GV Shiv the ashankar, and Caroline Uhler. Multi-domain translation between single-cell imaging and sequencing data using autoencoders. Nature Communications, 12(1):31, 2021.

[24] Taylor Rowe and Troy Day. The sampling distribution of the total correlation for multivariate gaussian random variables. Entropy, 21(10):921, 2019.

[25] Takuya Akiba, Shotaro Sano, Toshihiko Yanase, Takeru Ohta, and Masanori Koyama. Optuna: A next-generation hyperparameter optimization framework. In The 25th ACM SIGKDD International Conference on Knowledge Discovery & Data Mining, pages 2623–2631, 2019.

[26] Marija Milacic, Deidre Beavers, Patrick Conley, Chuqiao Gong, Marc Gillespie, Johannes Griss, Robin Haw, Bijay Jassal, Lisa Matthews, Bruce May, et al. The reactome pathway knowledgebase 2024. Nucleic Acids Research, 52(D1):D672–D678, 2024.

[27] Wolfgang Esser-Skala and Nikolaus Fortelny. Reliable interpretability of biology-inspired deep neural networks. NPJ Systems Biology and Applications, 9(1):50, 2023.

[28] David Antony Selby, Rashika Jakhmola, Maximilian Sprang, Gerrit Grossmann, Hind Raki, Niloofar Maani, Daria Pavliuk, Jan Ewald, and Sebastian J Vollmer. Visible neural networks for multi-omics integration: a critical review. bioRxiv, pages 2024–12, 2024.

[29] Emile Mathieu, Tom Rainforth, Nana Siddharth, and Yee Whye Teh. Disentangling disentanglement in variational autoencoders. In International Conference on Machine Learning, pages 4402–4412. PMLR, 2019.

[30] Benjamin Estermann and Roger Wattenhofer. Dava: Disentangling adversarial variational autoencoder. arXiv preprint arXiv:2303.01384, 2023.

[31] Ikram Eddahmani, Chi-Hieu Pham, Thibault Napoléon, Isabelle Badoc, Jean-Rassaire Fouefack, and Marwa El-Bouz. Unsupervised learning of disentangled representation via auto-encoding: A survey. Sensors, 23(4):2362, 2023.

[32] Scott Lundberg. A unified approach to interpreting model predictions. arXiv preprint arXiv:1705.07874, 2017.

[33] Avanti Shrikumar, Peyton Greenside, and Anshul Kundaje. Learning important features through propagating activation differences. In International Conference on Machine Learning, pages 3145–3153. PMlR, 2017.

[34] Marco Tulio Ribeiro, Sameer Singh, and Carlos Guestrin. " why should i trust you?" explaining the predictions of any classifier. In Proceedings of the 22nd ACM SIGKDD international conference on knowledge discovery and data mining, pages 1135–1144, 2016.

[35] Pascal Vincent, Hugo Larochelle, Yoshua Bengio, and Pierre-Antoine Manzagol. Extracting and composing robust features with denoising autoencoders. In Proceedings of the 25th international conference on Machine learning, pages 1096–1103, 2008.

[36] Johanna Zitt, Patrick Paitz, Fabian Walter, and Josefine Umlauft. Self-supervised coherence-based denoising on cryoseismological distributed acoustic sensing data. Authorea Preprints, 2024.

[37] Gökcen Eraslan, Lukas M Simon, Maria Mircea, Nikola S Mueller, and Fabian J Theis. Single-cell rna-seq denoising using a deep count autoencoder. Nature Communications, 10(1):390, 2019.

[38] Tianjiao Zhang, Hongfei Zhang, Jixiang Ren, Zhenao Wu, Zhongqian Zhao, and Guohua Wang. scdrmae: integrating masked autoencoder with residual attention networks to leverage omics feature dependencies for accurate cell clustering. Bioinformatics, 40(10):btae599, 2024.

[39] Jaesik Kim, Matei Ionita, Matthew Lee, Michelle L McKeague, Ajinkya Pattekar, Mark M Painter, Joost Wagenaar, Van Truong, Dylan T Norton, Divij Mathew, et al. Cytometry masked autoencoder: An accurate and interpretable automated immunophenotyper. bioRxiv, pages 2024–02, 2024.

[40] Maria Polychronidou, Jingyi Hou, M Madan Babu, Prisca Liberali, Ido Amit, Bart Deplancke, Galit Lahav, Shalev Itzkovitz, Matthias Mann, Julio Saez-Rodriguez, et al. Single-cell biology: what does the future hold?, 2023.

[41] Haotian Cui, Chloe Wang, Hassaan Maan, Kuan Pang, Fengning Luo, Nan Duan, and Bo Wang. scgpt: toward building a foundation model for single-cell multi-omics using generative ai. Nature Methods, pages 1–11, 2024.

[42] Qin Ma, Yi Jiang, Hao Cheng, and Dong Xu. Harnessing the deep learning power of foundation models in single-cell omics. Nature Reviews Molecular Cell Biology, pages 1–2, 2024.

[43] Minsheng Hao, Jing Gong, Xin Zeng, Chiming Liu, Yucheng Guo, Xingyi Cheng, Taifeng Wang, Jianzhu Ma, Xuegong Zhang, and Le Song. Large-scale foundation model on single-cell transcriptomics. Nature Methods, pages 1–11, 2024.

[44] F. Pedregosa, G. Varoquaux, A. Gramfort, V. Michel, B. Thirion, O. Grisel, M. Blondel, P. Prettenhofer, R. Weiss, V. Dubourg, J. Vanderplas, A. Passos, D. Cournapeau, M. Brucher, M. Perrot, and E. Duchesnay. Scikit-learn: Machine learning in Python. Journal of Machine Learning Research, 12:2825–2830, 2011.

[45] Pauli Virtanen, Ralf Gommers, Travis E. Oliphant, Matt Haberland, Tyler Reddy, David Cournapeau, Evgeni Burovski, Pearu Peterson, Warren Weckesser, Jonathan Bright, Stéfan J. van der Walt, Matthew Brett, Joshua Wilson, K. Jarrod Millman, Nikolay Mayorov, Andrew R. J. Nelson, Eric Jones, Robert Kern, Eric Larson, C J Carey, İlhan Polat, Yu Feng, Eric W. Moore, Jake VanderPlas, Denis Laxalde, Josef Perktold, Robert Cimrman, Ian Henriksen, E. A. Quintero, Charles R. Harris, Anne M. Archibald, Antônio H. Ribeiro, Fabian Pedregosa, Paul van Mulbregt, and SciPy 1.0 Contributors. SciPy 1.0: Fundamental Algorithms for Scientific Computing in Python. Nature Methods, 17:261–272, 2020.

[46] The scikit-learn-extra development team. scikit-learn-extra: a python module for machine learning that extends scikit-learn, 2020.

[47] Jianfang Liu, Tara Lichtenberg, Katherine A Hoadley, Laila M Poisson, Alexander J Lazar, Andrew D Cherniack, Albert J Kovatich, Christopher C Benz, Douglas A Levine, Adrian V Lee, et al. An integrated tcga pan-cancer clinical data resource to drive high-quality survival outcome analytics. Cell, 173(2):400–416, 2018.

[48] Jianjiong Gao, Bülent Arman Aksoy, Ugur Dogrusoz, Gideon Dresdner, Benjamin Gross, S Onur Sumer, Yichao Sun, Anders Jacobsen, Rileen Sinha, Erik Larsson, et al. Integrative analysis of complex cancer genomics and clinical profiles using the cbioportal. Science Signaling, 6(269):pl1–pl1, 2013.

[49] Kaiyi Zhu, Jaroslav Bendl, Samir Rahman, James M Vicari, Claire Coleman, Tereza Clarence, Ovaun Latouche, Nadejda M Tsankova, Aiqun Li, Kristen J Brennand, et al. Multiomic profiling of the developing human cerebral cortex at the single-cell level. Science Advances, 9(41):eadg3754, 2023.

[50] CZI Single-Cell Biology, Shibla Abdulla, Brian Aevermann, Pedro Assis, Seve Badajoz, Sidney M Bell, Emanuele Bezzi, Batuhan Cakir, Jim Chaffer, Signe Chambers, et al. Cz cellxgene discover: A single-cell data platform for scalable exploration, analysis and modeling of aggregated data. bioRxiv, pages 2023–10, 2023.

[51] Xuehua Ma, Zhiguang Zhao, Long Xiao, Weina Xu, Yahui Kou, Yanping Zhang, Gang Wu, Yangyang Wang, and Zhuo Du. A 4d single-cell protein atlas of transcription factors delineates spatiotemporal patterning during embryogenesis. Nature Methods, 18(8):893–902, 2021.

[52] François Chollet et al. Keras. https://keras.io, 2015.

[53] Chin-Wei Huang, Shawn Tan, Alexandre Lacoste, and Aaron C Courville. Improving explorability in variational inference with annealed variational objectives. Advances in Neural Information Processing Systems, 31, 2018.

[54] Hao Fu, Chunyuan Li, Xiaodong Liu, Jianfeng Gao, Asli Celikyilmaz, and Lawrence Carin. Cyclical annealing schedule: A simple approach to mitigating kl vanishing. arXiv preprint arXiv:1903.10145, 2019.

[55] Arek Kasprzyk. Biomart: driving a paradigm change in biological data management, 2011.

[56] Xiaojie Ma, Zhe Zhao, Lining Xiao, et al. A 4d single-cell protein atlas of transcription factors delineates spatiotemporal patterning during embryogenesis. Nature Methods, 18:893–902, 2021.

